# Capsaicin-induced endocytosis of endogenous presynaptic Ca_V_2.2 in DRG-spinal cord co-cultures inhibits presynaptic function

**DOI:** 10.1101/2022.10.03.510587

**Authors:** Krishma H Ramgoolam, Annette C Dolphin

## Abstract

The N-type calcium channel, Ca_V_2.2 is key to neurotransmission from the primary afferent terminals of dorsal root ganglion (DRG) neurons to their post-synaptic targets in the spinal cord. In this study we have utilized Ca_V_2.2_HA knock-in mice, because the exofacial epitope tag in Ca_V_2.2_HA enables accurate detection and localization of endogenous Ca_V_2.2. Ca_V_2.2_HA knock-in mice were used as a source of DRGs to exclusively study the presynaptic expression of N-type calcium channels in co-cultures between DRG neurons and wild-type spinal cord neurons. Ca_V_2.2_HA is strongly expressed on the cell surface, particularly in TRPV1-positive small and medium DRG neurons. Super-resolution images of the presynaptic terminals revealed an increase in Ca_V_2.2_HA expression and increased association with the post-synaptic marker Homer over time *in vitro*. Brief application of the TRPV1 agonist, capsaicin, resulted in a significant down-regulation of cell surface Ca_V_2.2_HA expression in DRG neuron somata. At their presynaptic terminals, capsaicin caused a reduction in Ca_V_2.2_HA proximity to and co-localization with the active zone marker RIM 1/2, as well as a lower contribution of N-type channels to single action potential-mediated Ca^2+^ influx. The mechanism of this down-regulation of Ca_V_2.2_HA involves a Rab 11a-dependent trafficking process, since dominant-negative Rab11a(S25N) occludes the effect of capsaicin on presynaptic Ca_V_2.2_HA expression, and also prevents the effect of capsaicin on action potential induced Ca^2+^ influx. Taken together, these data suggest that capsaicin causes a decrease in cell surface Ca_V_2.2_HA expression in DRG terminals via a Rab11a-dependent endosomal trafficking pathway.

## Introduction

Primary afferent axons of peripheral sensory dorsal root ganglion (DRG) neurons synapse in the dorsal horn of the spinal cord. The N-type calcium channel, Ca_V_2.2, is the main presynaptic voltage-gated calcium (Ca_V_) channel (VGCC) expressed in DRG neurons ^1^,^2^. It is a key mediator of nociceptive transmission, and it is important as a therapeutic target ^3^,^4^.

Ca_V_ channels are multi-subunit complexes composed of the central pore-forming α1 and auxiliary subunits, α_2_δ and β. A significant role for the α_2_δ-1 isoform has been established in chronic neuropathic pain ^5–8^. Furthermore, the gabapentinoids, including gabapentin, which are used in neuropathic pain conditions, act on α2δ-1 ^8,9^. The mechanism by which gabapentin reduces Ca^2+^ currents entails a reduction in trafficking of α_2_δ-1 and the associated calcium channel complex ^10^. This involves interference with recycling of α_2_δ-1 from Rab11a-positive recycling endosomes to the plasma membrane ^11,12^.

TRPV1 is a non-selective cation channel which is Ca^2+^-permeable and activated by the exogenous ligand capsaicin ^13^. It is located in distinct subpopulations of small and medium DRG neurons and plays an essential role in nociception, particularly noxious heat ^13–17^. Whereas brief exposure to capsaicin produces pain, it then desensitizes DRG neurons, resulting in reduced nociception ^18^, and a reduction in dorsal horn synaptic transmission ^19^. Ca^2+^ influx through TRPV1 channels initiates a cascade of events, one of them being changes in VGCC expression ^20^. However, the mechanisms behind the functional interaction between TRPV1 channels and VGCCs in primary nociceptors remains poorly understood.

Co-cultures between DRG and spinal cord neurons are an ideal model system for the study of synaptic transmission at primary afferent synapses ^21–24^. In previous studies, these co-cultures were also used to explore the mechanistic link between TRPV1 and Ca^2+^ signaling ^21^. It was demonstrated that a brief stimulation of TRPV1 with capsaicin resulted in glutamate release, independent of N-type calcium channels ^21^. It was also postulated that Ca^2+^ influx through TRPV1 mediated a Ca^2+^/calcineurin-dependent inhibition of N-type currents in DRG cell bodies ^20^.

In the present study, we examine the maturation of the localization of Ca_V_2.2 both in DRG neuronal somata and at their presynaptic terminals over time, by using DRG neurons from Ca_V_2.2_HA^KI/KI^ mice ^25^, co-cultured with wild-type spinal cord neurons. We then use these co-cultures to examine the effects of capsaicin on the distribution and presynaptic function of Ca_V_2.2_HA. We sought to determine whether capsaicin changes the expression of cell surface Ca_V_2.2, and to understand the mechanisms influencing presynaptic localization and function of this channel. Our results reveal a dramatic effect of brief capsaicin exposure on Ca_V_2.2 distribution in the somata and in presynaptic terminals of DRG neurons, which occurs with a slow time course and involves a Rab11a-dependent pathway. Our results support the view that presynaptic Ca_V_2.2 channels are highly dynamic in their localization and activity-dependent turnover.

## Materials and Methods

### Animals

The Ca_V_2.2_HA^KI/KI^ C57BL/6 mouse line described previously ^25^ and wild-type C57BL/6 mice were housed in groups of no more than five on a 12-h:12-h light: dark cycle; food and water were available ad libitum. Both Ca_V_2.2_HA^KI/KI^ and Ca_V_2.2^WT/WT^ mice were obtained by breeding from homozygotes. All experimental procedures were covered by UK Home Office licenses, had local ethical approval by University College London (UCL) Bloomsbury Animal Welfare and Ethical Review Body. All cultures were prepared from P0-P1 male and female mice.

### DRG-spinal cord neuronal co-cultures

DRG neurons from Ca_V_2.2_HA^KI/KI^ mice were cultured with spinal cord neurons from Ca_V_2.2^WT/WT^ P0/1 mice. DRG and spinal cords were extracted in ice cold dissection medium (Leibovitz’s L15 Medium without supplements; Gibco). The spinal cord was dissected in small segments (~0.5 mm) and then digested in 2.5 % trypsin (Gibco)and 1200 U/μl DNase I (Sigma Aldrich) for 23 min at 37°C in a 5 % CO2 incubator. The spinal cord sections were then washed with 37°C culture medium I (10 % fetal bovine serum, 1 unit/ml penicillin, 1μg/ml mg streptomycin and DMEM; Gibco). Following this, the spinal cord was triturated twice gently with fire polished glass pipettes. Spinal cord suspensions were plated onto either 22 or 25 mm glass coverslips which were coated with poly-L-lysine and laminin.

DRGs were dissected and incubated in Hanks’ Balanced Salt solution (HBSS; Gibco^TM^) and kept at 4°C until all DRG tissue was removed. The DRGs were incubated in DRG enzyme solution (1000 U/ml DNAse I, 3.75 mg/ml dispase and 0.8 mg/ml collagenase type 1A; Gibco) for 21 min at 37°C in a 5 % CO_2_ incubator. Culture medium I was added to the digested tissue, which was then transferred to a 1.5ml Eppendorf tube and centrifuged at 1000 rpm for 5 min. The supernatant was removed, the pellet was re-suspended in culture medium I and triturated three times with fire-polished glass pipettes to produce a single cell suspension. The cell suspension was centrifuged at 1000 rpm for 5 min and supernatant removed. Culture medium I was added to the pellet before the DRG neurons were plated on to the spinal cord neurons. The dishes were flooded with culture medium I (at 37°C), 1 h after plating. In this study, co-cultures used at DIV 1, 7 and 14 are referred to as immature cultures. Co-cultures utilized at DIV 21 and 28 are classified as mature cultures.

For experiments involving transfection, DRG neurons were transfected before plating. To ensure sufficient transfection of DRG neurons, DRG tissue was collected from three mice (P0/P1). After dissociation, the DRG cell suspension was washed with 1 ml of 37°C HBSS and centrifuged at 1000 rpm for 5 min. The supernatant was discarded, cell pellet was resuspended in 100 ml Nucleofector™ (Rat Neuron Nucleofector kit, Lonza) transfection reagent and electroporated with 2 μg of cDNA mix according to the manufacturer’s protocol. The following mixes were transfected separately for the experiments in this study: synaptophysin-GCaMP6f (Sy-GCaMP6f) and VAMP-mOrange 2 (VAMP-mOr2; ^26^) (3:1); empty vector and mCherry (1:1); Rab11a(S25N) and mCherry (1:1); Empty Vector and mCherry (1:1); Rab11a(S25N), Sy-GCaMP6f, VAMP-mOr2 (1:1:1) and Empty Vector, Sy-GCaMP6f, VAMP-mOr2 (1:1:1). For expression in DRG neurons, Rab11a(S25N) ^12^ was subcloned into the vector pCAGGs. Electroporated cells were then incubated in 1 0% FBS-RPMI (Roswell Park Memorial Institute medium) for 5 min at 37°C in a 5 % CO_2_ incubator. The DRG cell suspension was finally added dropwise on top of the plated spinal cord neurons.

For both electroporated and non-electroporated cells, culture medium I was added to the cells 1 h after plating. The following day, culture medium I was replaced with culture medium II (DMEM supplemented with 24 μg/ml insulin (Sigma Aldrich), 100 μg/ml transferrin (EMD Millipore), 5 % horse serum, 2X B27 supplement (Gibco), 1X GlutaMAX™, 0.5 ng/ml NGF (Sigma Aldrich) and 1 unit/ml penicillin, 1μg/ml streptomycin). 50 % of the growth medium was replaced every 3 - 4 days with fresh culture medium II. To inhibit the proliferation of non-neuronal cells, after 48 h, 5 μM cytosine-A-D-arabinofuranoside (AraC; Gibco) was added to cultures for 12 h. Two days later, cells were again treated with 5 μM AraC for 12 h.

### Immunolabelling of DRG - spinal cord neuronal co-cultures

All immunocytochemistry protocols included the use of blocking buffer (20 % horse serum in PBS), antibodies were diluted in antibody solution (10 % horse serum in PBS) and for permeabilized conditions 0.1-0.3 % v/v Triton-X 100 in PBS was included. All co-cultures were fixed and post-fixed (before and after incubation with primary rat anti-HA antibody (Roche), respectively) using 4 % paraformaldehyde (PFA) and 4 % sucrose in PBS for 5 min at room temperature (20°C). All primary and secondary antibodies were applied separately and consecutively to one another to prevent problems with cross-reactivity of antibodies.

#### Cell surface Ca_V_2.2_HA in DRG cell bodies

Co-cultures were fixed and blocked for a minimum of 1 h at 20°C. To prevent non-specific binding of secondary antibodies, cultures were first incubated with rat anti-HA (1:100) antibody overnight at 4°C followed by donkey anti-rat Alexa Fluor (AF) 488 (1:500; Invitrogen) for 1.5 h at 20°C before staining for other markers. Unbound primary antibody was washed off using PBS and cells were post-fixed.

#### Ca_V_2.2_HA, Homer and vGluT2

Co-cultures were fixed and blocked for 1 h at 20°C. First, for Ca_V_2.2_HA immunolabelling, cultures were incubated in antibodies as described above. Next, cultures were permeabilised by incubation in antibody solution containing 0.3 % Triton X-100. Cultures were incubated with rabbit anti-Homer antibody (1:2000; Frontier Institute) antibody overnight at 4°C. Subsequently, cultures were incubated with donkey anti-rabbit AF 633 antibody (1:500; Invitrogen) for 1.5 h at 20°C. Finally, for vGluT2 immunolabelling, neurons were incubated with guinea pig (GP) anti-vGluT2 (1:5000; Merk Millipore) antibody overnight at 4°C. The next day, secondary donkey anti-GP AF 594 (1:500; Invitrogen) antibody was applied to neurons for 1.5 h at 20°C.

#### Ca_V_2.2_HA and RIM 1/2

Co-cultures were fixed and blocked for 1 h at 20°C. First, for Ca_V_2.2_HA immunolabelling, cultures were incubated using the protocol above. Following this, neurons were incubated with rabbit-anti-RIM 1/2 (1:200; Synaptic Systems) antibody overnight at 4°C. Next, secondary donkey anti-rabbit AF 594 (1:500; Invitrogen) antibody was applied to neurons for 1.5 h at 20°C.

#### Capsaicin application and immunocytochemical protocol

1 μM capsaicin diluted in Krebs-Ringer-Hepes (KRH) was applied to co-cultured neurons at 21 DIV for 2 min at 37°C. For controls, neurons were incubated similarly in KRH but without capsaicin. After incubation with capsaicin, cultures were briefly washed and incubated in culture medium II for a 0-, 20-, 40- or 60-min rest period at 37°C. Co-cultures were then fixed at 20°C for 5 min. Following this, cells were blocked for 1 h at 20°C.

#### Ca_V_2.2_HA and TRPV1

Co-cultures were fixed and blocked for 1 h at 20°C. First, for Ca_V_2.2_HA immunolabelling, cultures were incubated in antibodies as described above. After this, for TRPV1 immunolabelling, neurons were incubated with goat anti-TRPV1 (1:500; Santa Biotech) antibody overnight at 4°C in permeabilised conditions (0.1 % Triton X-100 in antibody solution), followed by donkey anti-goat AF488 antibody (1:500; Invitrogen) for 1.5 h at 20°C.

#### Ca_V_2.2_HA, TRPV1 and RIM 1/2

For cell surface Ca_V_2.2_HA and TRPV1 labelling, the protocol above was used. Following this, cultures were incubated in rabbit anti-RIM 1/2 antibody (1:200) followed by donkey anti-rabbit AF647 antibody (1:500; Invitrogen).

#### Ca_V_2.2_HA, RIM 1/2 and mCherry

For Ca_V_2.2_HA and RIM 1/2 immunolabelling, the protocols above were used. Subsequently, co-cultures were incubated in GP anti-red fluorescent protein (RFP) antibody at 1:500 (Synaptic Systems) overnight at 4°C. Next, secondary donkey anti-GP AF 594 antibody was applied to coverslips at 1:500 (Invitrogen) for 1.5 h at 20°C.

Following all immunocytochemical protocols, nuclei were stained with DAPI (4’,6-diamidino-2-phenylindole) (500 nM) before mounting on slides using Vectashield (Vector Laboratories) to reduce photobleaching.

### Image acquisition and analysis

Co-cultures were examined using super resolution Airyscan mode imaging on a LSM 780 confocal microscope (Zeiss) with x 63 objective (1768 × 1768 pixels) as z-stacks (0.173 μm optical section). Superresolution images then underwent pixel reassignment and Airyscan processing (6x) using Zen software. Images were acquired with constant settings in each experiment from at least 3 separate cultures per experiment.

To quantify Ca_V_2.2_HA density and its association with vGlut2, Homer and RIM 1/2, the Image J plugin: Distance Analysis (DiAna) ^27^ was used. DiAna was used to quantify the distance measurements between centres of co-localized objects, percentages of co-localizing volumes for each object’s pair and mean intensity of puncta.

For cell body analysis, using ImageJ software (Schneider et al., 2012), every DRG neuron with a visible nucleus with Ca_V_2.2_HA immunolabelling was measured ^25^. A 10-pixel wide line (0.9μm) was drawn following the perimeter of the cell from which the perimeter length was recorded as an estimation of the size of the cell (small <61 μm, medium 61-94 μm or large >94 μm;^28^), and the mean membrane HA fluorescence intensity.

### Live cell Ca^2+^ imaging

#### Ca^2+^ imaging of co-cultures

Live cell Ca^2+^ imaging was performed at DIV 7, 14, 21 and 28, as previously described with minor modifications ^26,29^. Coverslips were mounted in a rapid-switching, laminar-flow perfusion, and stimulation chamber (RC-21BRFS, Warner Instruments) on the stage of an epifluorescence microscope (Axiovert 200 M, Zeiss). Live cell images were acquired with an Andor iXon+ (model DU-897U-CS0-BV) back-illuminated EMCCD camera using OptoMorph software (Cairn Research, UK). White and 470 nm LEDs served as light sources (Cairn Research, UK). Fluorescence excitation and collection was done through a Zeiss 40 × 1.3 NA Fluor objective using 450/50 nm excitation and 510/50 nm emission and 480 nm dichroic filters (for sy-GCaMP6f) and a 545/25 nm excitation and 605/70 nm emission and 565 nm dichroic filters (for mOrange2). Action potentials (AP) were evoked by passing 1 ms current pulses via platinum electrodes. Cells were perfused (0.5 ml.min^-1^) in a saline solution at 25°C containing (in mM) 119 NaCl, 2.5 KCl, 2 CaCl_2_, 2 MgCl_2_, 25 HEPES (buffered to pH 7.4), 30 glucose, 10 μM 6-cyano-7-nitroquinoxaline-2,3-dione (CNQX, Sigma) and 50 μM D,L-2-amino-5-phosphonovaleric acid (AP5, Sigma). Images were acquired at 100 Hz over a 512 × 266 pixel area in frame transfer mode (exposure time 7 ms) and analyzed in ImageJ (http://rsb.info.nih.gov/ij) using a custom-written plugin (http://rsb.info.nih.gov/ij/plugins/time-series.html). Successfully transfected neurons were identified by visualizing sy-GCaMP6f fluorescence in response to a 33 Hz stimulation for 180 ms every 4 s. Subsequently, single stimulations of 1 ms (mimicking a single AP) were repeated 5 times with 30 s intervals. Regions of interest (ROI, 2 μm diameter circles) were placed around synaptic boutons responding to an electrical stimulation of 10 AP at 60 Hz. Functional synaptic boutons were identified by the increase of fluorescence of VAMP-mOr2 in response to 200 AP stimulation at 10 Hz (in this case images were acquired at 2 Hz with 50 ms exposure time). ω-conotoxin GVIA (1 μM; CTX, Alomone Labs) was perfused for at least 2 min. To determine if any of the observed reduction in fluorescence was due to bleaching, control experiments were performed where CTX was replaced with normal imaging medium and perfused for 2min. Cells were then re-stimulated and a reduction of 6.9 ± 4.0 % was recorded during 1 AP stimulation. All the values shown here have been adjusted for this reduction.

#### Ca^2+^ imaging of capsaicin treated co-cultures following perfusion of control medium for 20 - 60-min rest period

An initial protocol consisting of 1 AP and 200 APs stimulations was applied to assess the baseline responses. After 5 min, either capsaicin or control medium was applied for 2 min. Control medium was then perfused for 20, 40, or 60 min. To assess the contribution of N-type calcium channels to the Ca^2+^transient following 1AP, CTX was applied for 2 min. A final 1 AP train stimulation was applied to the cultures and the initial Ca^2+^ transients were used to normalise the final Ca^2+^ transient to determine the contribution of N-type VGCCs. To determine if any of the observed reduction in fluorescence of capsaicin-treated or control-treated neurons was due to bleaching, control experiments were performed where CTX was replaced with normal imaging medium and perfused for 2 min. Following this, cells were perfused in normal live imaging medium for 20, 40 or 60 min. Control-treated cells were then re-stimulated with 1 AP and a reduction of 4.8 ± 7.0 %, 11 ± 3.0 % and 5 ± 2% was recorded for cells perfused for 20, 40 or 60 min, respectively. Capsaicin treated cells were also re-stimulated with 1 AP and a reduction of 0.03 ± 0.01 %, 0.3 ± 0.03 % and 1.8 ± 0.6 % was recorded for cells perfused for 20, 40 or 60 min, respectively. Values shown in this study have been adjusted for these reductions.

### Statistical analysis

Data were analyzed with Prism 9.0 (GraphPad Software). Where error bars are shown, they are SEM; *“n”* refers to the number of separate cultures used (termed experiments), unless indicated otherwise. Statistical significance between two groups was assessed by Student’s *t* test, as stated. One-way ANOVA and stated post hoc analysis, recommended as appropriate by Prism, were used for comparison of means between three or more groups.

## Results

### Ca_V_2.2_HA expression in DRG - spinal cord neuron co-cultures

In the present study, a co-culture system was devised to recapitulate major characteristics of peripheral neuron maturation, synapse formation and function. We combined DRG neurons cultured from Ca_V_2.2_HA^KI/KI^ mice with spinal cord neurons from Ca_V_2.2^WT/WT^ mice (Fig. 1A). This ensured that the changes in Ca_V_2.2_HA being investigated were only those occurring in the presynaptic DRG neurons and their terminals. DRG glial cells and spinal cord astrocytes served as substrates for the co-cultured neurons. DRG neurites have shown a preferential growth *in vitro* into dorsal spinal cord explants ^30^. The time course for functional synapse formation between DRG neurons with their dorsal horn partners has been reported to commence at day *in vitro* (DIV) 5 ^31^, and previous studies have investigated these processes in cocultures for up to 4 weeks ^22,23^.

**Figure 1:**
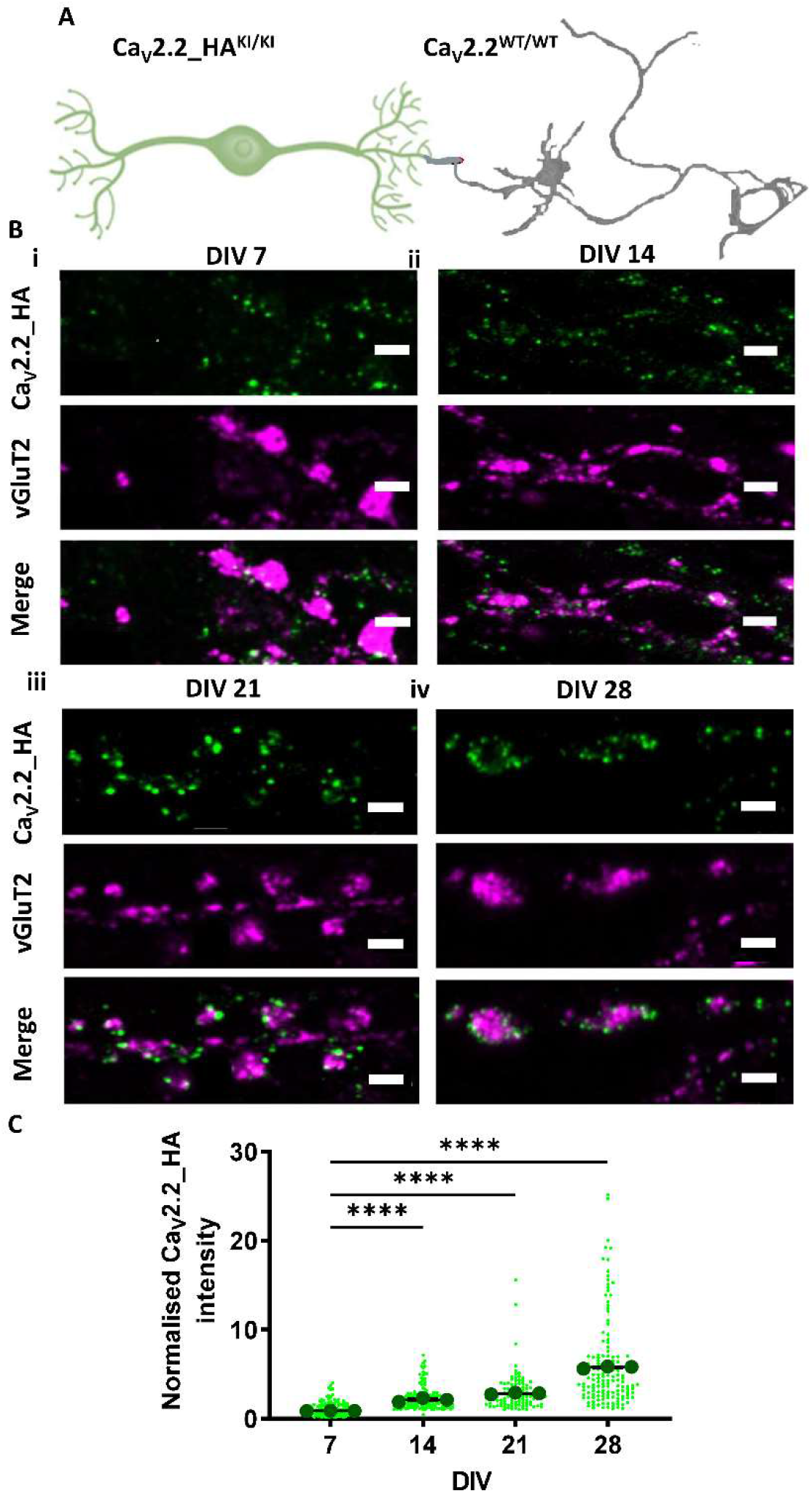
The development of Ca_V_2.2_HA presynaptic expression in DRG-spinal cord neuron co-cultures. (**A**) Schematic diagram of co-cultures composed of DRG neurons from Ca_V_2.2_HA^KI/KI^ mice (green) cultured with spinal cord neurons from Ca_V_2.2^WT/WT^ mice (grey). (**B**) Airyscan images of co-cultures at (**i**) DIV 7, (**ii**) DIV 14, (**iii**) DIV 21 and (**iv**) DIV 28. Ca_V_2.2_HA (top panel; green), vGluT2 (middle panel; magenta) and merged image (bottom panel). Scale bars: 2 μm. (**C**) Ca_V_2.2_HA intensity measured at presynaptic terminals of DRG neurons from co-cultures fixed at DIV 7, 14, 21 and 28 (green points). Individual data points represent 150 vGlut2 positive puncta also positive for Ca_V_2.2_HA. Mean ± SEM of the three experiments (black circles) are superimposed. Mean Ca_V_2.2_HA intensity of each experiment was normalised to that at DIV 7. Statistical analysis: one-way ANOVA with Sidak’s multiple selected comparison post-hoc test; **** p<0.0001.

The clustering of synaptic vesicles at presynaptic active zones is one of the initial signs of synapse formation. All primary afferents use glutamate as their major neurotransmitter, thus having an excitatory effect on their postsynaptic targets ^32,33^. Spinal cord synaptic vesicles are enriched in vesicular glutamate transporters (vGluT)-1 and vGluT2 ^34,35^. vGluT2 immunolabelling has been shown to be most prominent in Lamina II of the spinal cord, suggesting that it is associated with glutamatergic transmission from small diameter nociceptors ^34^. For this reason, vGluT2 was used here as a presynaptic marker to identify glutamatergic synapses.

Ca_V_2.2_HA expression was examined in co-cultures at different timepoints: DIV 7, 14, 21 and 28 (Figs. 1B, C). At DIV 7, Ca_V_2.2_HA puncta were observed diffusely throughout the neurites, and were not strongly associated with vGlut2 puncta (Figs. 1Bi, C). In contrast, in DIV 28 co-cultures, both Ca_V_2.2_HA and vGlut2 showed a strong punctate staining profile along the processes (Fig. 1 Biv). Ca_V_2.2_HA was distributed around a central core of vGlut2 more frequently at DIV 28 compared to DIV 7 (Figs. 1Bi and iv). Furthermore, the intensity of Ca_V_2.2_HA puncta, co-localized to vGluT2, normalised to the average intensity measured at DIV 7, was significantly higher at DIV 28 (5.9 ± 0.1) than at DIV 7 (1.0 ± 0.1) (Fig. 1C). A gradual increase in Ca_V_2.2_HA intensity was also observed at the intermediate timepoints DIV 14 (2.3 ± 0.1) and DIV 21 (3.0 ± 0.1) (Figs. 1Bii, Biii and C).

### Development of Ca_V_2.2_HA expression at presynaptic boutons of DRG - spinal cord neuron co-cultures

Presynaptic Ca_V_2.2_HA was further examined using a panel of markers to study the temporal pattern of synapse formation between DRG and spinal cord neurons (Fig. 2 and Supplementary Fig. 1). We assessed the additional co-localization of Ca_V_2.2_HA with RIM 1/2 and Homer. Since Homer immunostaining reveals the majority of excitatory synapses ^36^, boutons were chosen based on the associated presence of Homer immunoreactivity. Furthermore, synaptic Ca_V_2.2 channels should be localized adjacent to docked and primed synaptic vesicles, and the presynaptic active zone was identified by the presence of RIM 1/2. At DIV 7, weak Ca_V_2.2_HA immunolabelling, associated with these synaptic markers, could be seen in immature cultures (Figs. 2Ai and Aii). In contrast, in mature DIV 28 cultures, Ca_V_2.2_HA-positive puncta appeared clearly defined in close apposition to Homer, vGluT2 and RIM 1/2 puncta (Figs. 2Ai and Aii). The appearance of intense punctate staining along the axonal processes was first seen at DIV 21 (Supplementary Fig. 1). In addition to elongated punctate structures, Ca_V_2.2_HA can also be seen distributed around a central core of vGluT2 associated with Homer, resembling glomerular synapses ^37^. The rosette-shaped clusters of Ca_V_2.2_HA were comprised of up to five puncta (Supplementary Figs. 1A, B). These patterns of immunoreactivity are consistent with our *in vivo* study using Ca_V_2.2_HA^KI/KI^ mice ^25^. There was no difference between the relative distance between Ca_V_2.2_HA and vGluT2 puncta at DIV 7 (0.36 ± 0.02 μm) and DIV 28 (0.35 ± 0.02 μm) (Fig. 2Bi), or between Ca_V_2.2_HA and RIM 1/2 puncta at DIV 7 (0.31 ± 0.01 μm) and DIV 28 (0.29 ± 0.01 μm) (Fig. 2Ci). However, a significant decrease was observed in the distance between Ca_V_2.2_HA and Homer at DIV 28 (0.24 ± 0.01 μm) compared to DIV 7 (0.38 ± 0.01 μm) (Fig. 2Di), indicative of closer association with post-synaptic densities. Additionally, the overall percentage co-localization between Ca_V_2.2_HA and vGluT2, Homer and RIM 1/2 increased by 22.6, 14.0, and 21.3%, respectively, when comparing DIV 7 to DIV 28 (Figs. 2Bii, Cii and Dii). This was also seen at the intermediate time point DIV 21 (Supplementary Figs. 1C, D and E). Taken together, these data suggest that the expression pattern of presynaptic Ca_V_2.2_HA in these co-cultures is developmentally regulated, consistent with the formation of mature synapses between DIV 21 and DIV 28.

**Figure 2:**
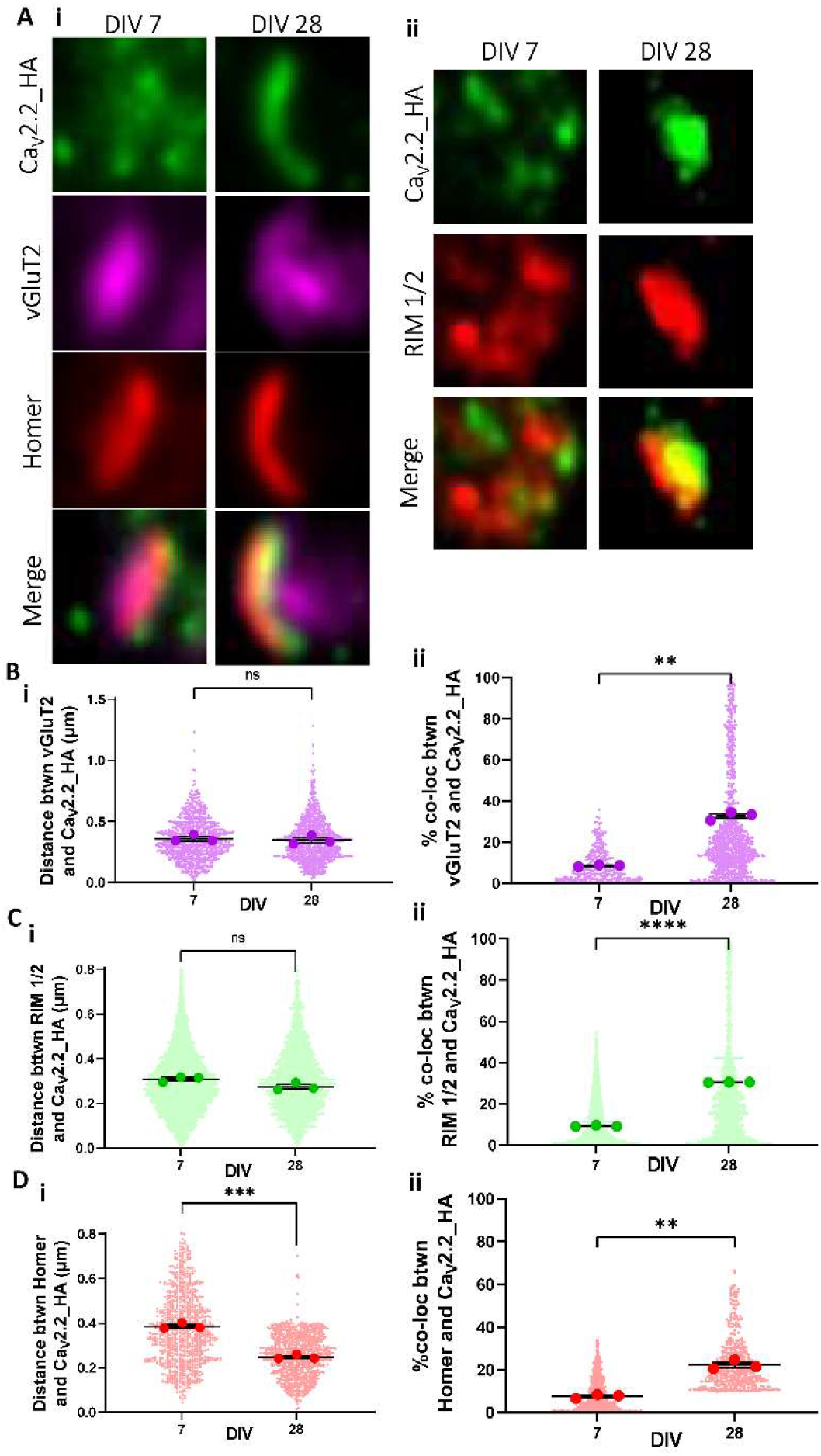
Increased co-localization of Ca_V_2.2_HA with vGluT2, Homer and RM 1/2 at the presynaptic membrane in DRG-spinal cord co-cultures at DIV 28. (**A**) Representative images of single optical sections of synaptic puncta, (2 x 2 μm ROI), for DIV 7 (left) and DIV 28 (right) showing (**i**) Ca_V_2.2_HA (green), vGluT2 (magenta), Homer (red) and merged panels and (**ii**) Ca_V_2.2_HA (green), RIM 1/2 (red) and merged panels. Distance measurements between centres of co-localized objects (**Bi**) Ca_V_2.2_HA and vGluT2, (**Ci**) Ca_V_2.2_HA and RIM 1/2 and (**Di**) Ca_V_2.2_HA and Homer. Measurements of the percentage colocalizing volume for each object’s pair (**Bii**) Ca_V_2.2_HA and vGluT2, (**Cii**) Ca_V_2.2_HA and RIM 1/2 and (**Dii**) Ca_V_2.2_HA and Homer. Individual data points represent distance and percentage measurements between co-localizing objects, 900, 1400 and1300 for Ca_V_2.2_HA and vGluT2, Ca_V_2.2_HA and RIM 1/2 and Ca_V_2.2_HA and Homer, respectively from DIV 7 and 28. Mean of each experiment is shown with larger symbols. Mean ± SEM of the three experiments is superimposed. Statistical analysis: unpaired t test with Welch’s correction; ****p<0.0001, ***p<0.001, **p<0.01, ns = not significant.

### Development of functional synapses dependent on N-type channels in DRG - spinal cord neuron co-cultures

After characterization of the developmental organization of endogenous Ca_V_2.2_HA at DRG presynaptic terminals within co-cultures, we next sought to establish whether these synaptic boutons were functional using Ca^2+^ imaging at increasing DIV. DRG neurons were transfected, prior to plating, with the functional presynaptic reporter synaptophysin coupled to the genetically encoded Ca^2+^ indicator GCaMP6f (sy-GCaMP6f) ^26^, and with a reporter of presynaptic exocytosis, vesicle associated membrane protein (VAMP) tagged with the pH-sensitive fluorescent protein mOrange 2 (VAMP-mOr2) (Fig. 3A). To assess local Ca^2+^transients, in response to a single action potential (AP), a train of 1 AP stimuli was applied (Fig. 3B). An increase of VAMP-mOr2 fluorescence in response to a stimulus of 200 APs at 10 Hz was then used to identify the responses from functional synaptic boutons, as previously described ^29,38^ (Figs. 3A, C). Additionally, to identify the contribution of N-type VGCCs to the Ca^2+^ transients, we used the specific Ca_V_2.2 inhibitor, ω-conotoxin GVIA (CTX). Application of CTX to co-cultures at DIV 7 and 28 indicated that N-type VGCCs mediated 76.6 ± 5.1 % and 73.9 ± 4.3 % of Ca^2+^ entry due to 1 AP, respectively (Fig. 3D), with similar results at intermediate DIVs (Supplementary Fig. 2A). However, there was a marked increase in the percentage of functional synaptic boutons at DIV 28 (30.2 ± 3.1 %) compared to DIV 7 (13.4 ±1.9 %) (Fig. 3E). A similar increase was observed at DIV 21 (Supplementary Fig. 2B). These results suggest that DRG and spinal cord neurons readily form functional synapses with increasing time in co-culture, from DIV 7 to 28 and that N-type calcium channels are responsible for a large proportion of the Ca^2+^transient seen following 1 AP stimulation.

**Figure 3:**
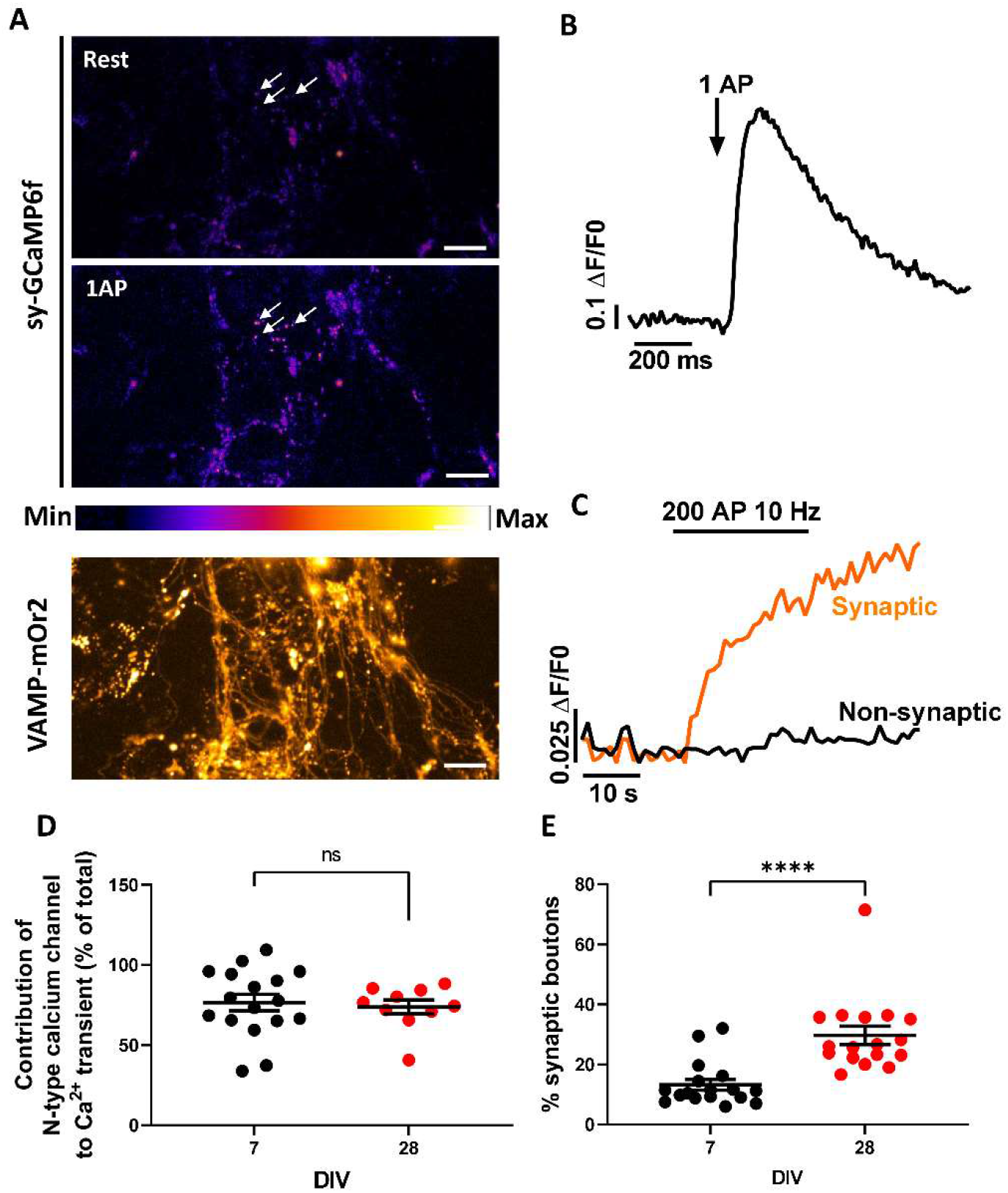
Increase in presynaptic Ca^2+^ transients associated with synaptic boutons relative to non-releasing boutons in co-cultures between DIV 7 and 28. (**A**) GCaMP6f fluorescence changes in presynaptic terminals of DRG neurons expressing sy-GCaMP6f and VAMP-mOr2, in response to electrical field stimulation. Sy-GCaMP6f fluorescence: at rest (top) and after 1AP (middle) stimulation. White arrows point to examples of transfected boutons. The pseudocolour scale is shown below the second panel. The bottom panel shows presynaptic terminals expressing VAMP-mOr2 following 200 AP stimulation. Scale bar: 10 μm. (**B**) Example of sy-GCaMP6f fluorescence (Ca^2+^ transient) increase in response to 1AP stimulation in DRG neuron terminals (averaged trace from 5 neurons). (**C**) Examples of VAMP-mOr2 fluorescence in response to 200 APs at 10 Hz from DRG neuron terminals, used to identify vesicular release from presynaptic boutons: each individual bouton was analysed and grouped into “non-synaptic” (black trace) or “synaptic” (orange trace; positive response to 200 AP at 10 Hz) groups depending on whether there was no variation or an increase in fluorescence was recorded in response to stimulation. The traces correspond to the average responses from 50 boutons each. (**D**) Contribution of N-type VGCCs to the Ca^2+^ transient in response to 1AP; 76.6 ± 5.1 % (n = 17) and 73.9 ± 4.3 % (n = 10) for DIV 7 (black circles) and 28 (red circles), respectively (measured from synaptic boutons). (**E**) % of synaptic-positive boutons at DIV 7 (black circles) and 28 (red circles), n = 17 for both DIV 7 and 28. n number refers to the number of experiments at each timepoint *in vitro*. Statistical analysis: Student’s t test with Welch’s correction; **** p<0.0001 and ns = not significant.

### Cell surface Ca_V_2.2_HA is preferentially expressed in TRPV1-positive DRG neurons in co-cultures

Ca_V_2.2_HA distribution was also compared in DRG neuronal somata with respect to cell size and DIV (Supplementary Figs. 3Ai and ii). A ring-like pattern of Ca_V_2.2_HA immunolabelling at the cell surface is observed in both small and medium DRG neurons which diminished with increasing time in culture from DIV 7 to DIV 21 - 28 (Supplementary Figs. 3A - C).

Capsaicin sensitivity is an important pharmacological trait of a major subset of nociceptive sensory neurons which mediates its effects through TRPV1 ^13,39^. As described above, capsaicin has been reported to profoundly inhibit VGCC currents in DRG neurons ^40^. To understand the association between TRPV1 and Ca_V_2.2 in DRG populations, co-cultures were immunolabelled for both Ca_V_2.2_HA and TRPV1. There was a clear ring of Ca_V_2.2_HA at the plasma membrane of most neurons positive for TRPV1, and this was absent from Ca_V_2.2 wild-type neurons (Fig. 4A). Although cell surface Ca_V_2.2_HA immunoreactivity can also be seen, to a lesser extent, on TRPV1-negative neurons (Fig. 4A), 66.4 ± 6.3 % of Ca_V_2.2_HA-positive neurons expressed TRPV1 (Fig. 4B). TRPV1 expression was found to be highest in small and medium DRG neurons, as also seen in previous studies ^16,41^. Cell surface Ca_V_2.2_HA expression levels were higher, particularly in medium TRPV1-positive neurons, compared to their TRPV1-negative counterparts (Fig. 4C). These data suggest that Ca_V_2.2_HA is preferentially expressed in TRPV1-positive medium DRG neurons.

**Figure 4:**
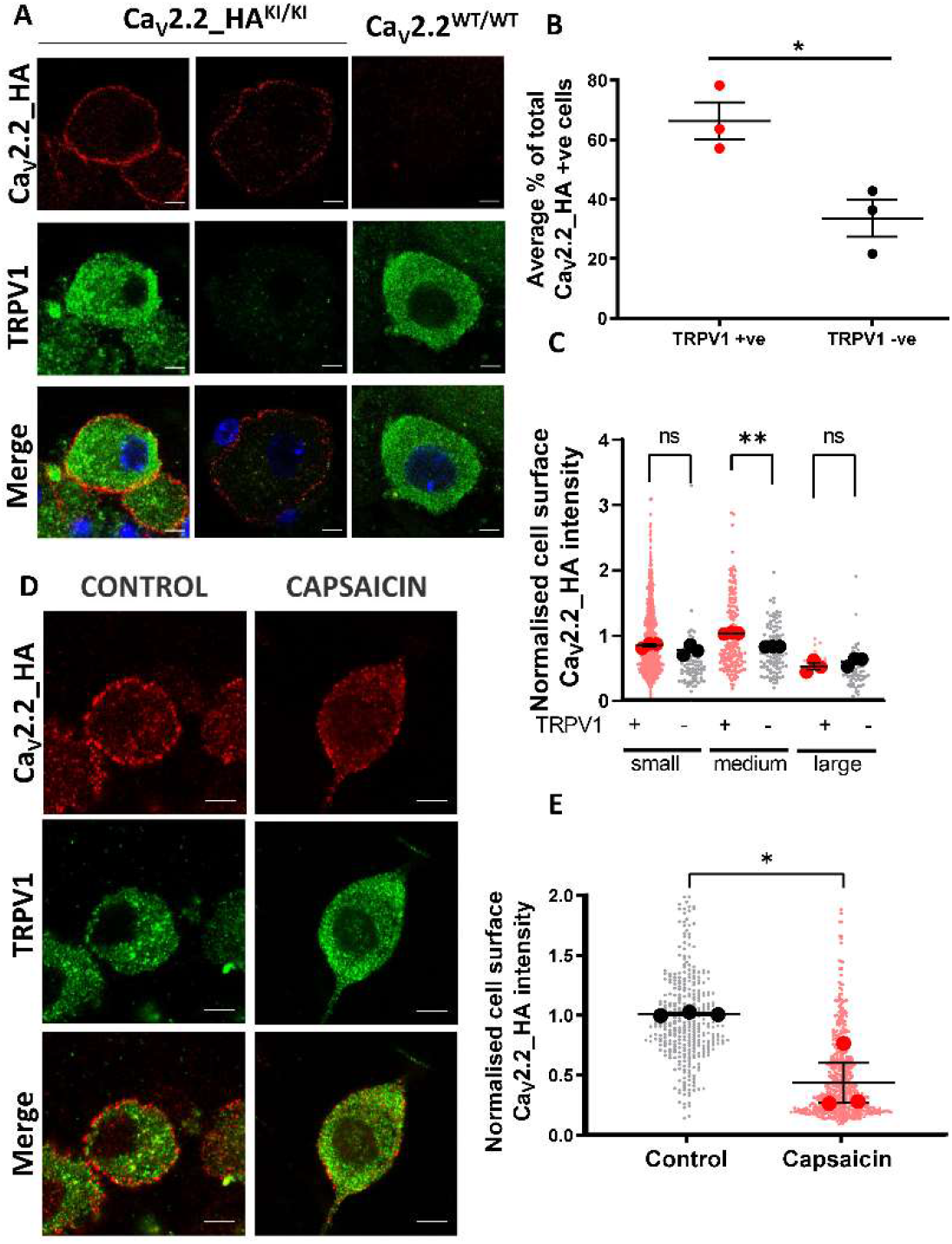
Decrease in cell surface Ca_V_2.2_HA in small DRG neurons following incubation with capsaicin. (**A**) Images of Ca_V_2.2_HA^KI/KI^ and Ca_V_2.2^WT/WT^ DRG neurons in DIV 21 co-cultures, showing Ca_V_2.2_HA staining before permeabilization (top; red), TRPV1 staining following permeabilization (middle, green) and merged images (bottom), for representative Ca_V_2.2_HA^KI/KI^ (left and middle panels) and Ca_V_2.2^WT/WT^ (right panel) DRG neurons. Scale bars 5μm. (**B**)Quantification of the percentage of cells with cell-surface Ca_V_2.2_HA that were either positive (red circles) or negative (black circles) for TRPV1. Individual data points represent the mean data from three separate experiments and a total of 311 DRG neurons. *p < 0.05 (Student’s t test). (**C**) Normalised cell surface Ca_V_2.2_HA intensity with respect to cell size: small, medium and large DRG neurons that are either TRPV1-positive (red circles) or TRPV1-negative (black circles). Individual data points represent normalised Ca_V_2.2_HA intensity measured from all Ca_V_2.2_HA positive cells from three separate experiments (with mean of each shown in larger circles) and a total of 185, 77, 184, 102, 14 and 63 DRG neurons, respectively. Mean ± SEM of the three experiments is superimposed. Statistical analysis: one-way ANOVA with Sidak’s multiple selected comparison post hoc test; **p<0.01, ns, not significant. (**D**) Capsaicin or control medium applied to DIV 21 co-cultures for 2 min, followed by 60 min rest at 37°C prior to fixation. Representative small DRG neurons in control (left) and capsaicin (right) conditions, showing Ca_V_2.2_HA (top; red), TRPV1 (middle; green) and merged image (bottom). Scale bars: 5 μm. (**E**) Normalized cell surface Ca_V_2.2_HA intensity measured from control or capsaicin-treated small DRG neurons following 60 min rest at 37°C. Individual data points represent normalized Ca_V_2.2_HA intensity measured from three separate experiments and a total of 357 and 546 small DRG neurons from control (black circles) and capsaicin conditions (red circles). Mean of each experiment shown in larger symbols. Mean ± SEM (n=3) is superimposed. Statistical analysis: Student’s t-test * p<0.05.

### Capsaicin reduces somatic cell surface expression of Ca_V_2.2_HA

To study the effect of capsaicin on cell surface Ca_V_2.2_HA expression and its time course, co-cultures were incubated with either 1 μM capsaicin or control buffer solution for 2 min at 37°C and then allowed to rest at 37°C for 0, 20, 40 or 60 min. Capsaicin (2 min) produced a dramatic reduction, by 57 % in Ca_V_2.2_HA cell surface labelling in small DRG neurons after 60 min (Figs. 4D, E) and after 40 min rest (Supplementary Fig. 4). Moreover, a similar pattern was observed in medium DRG neurons (Supplementary Fig. 5). In contrast, in controls we observed no change in plasma membrane labelling of Ca_V_2.2_HA in either small or medium DRG neurons at any time point (Supplementary Figs. 4A, C and 5A, C). These data indicate that capsaicin induces a decrease in cell surface Ca_V_2.2_HA expression in both small and medium DRG neurons, which occurs with a slow time-course.

### Capsaicin modulates presynaptic Ca_V_2.2_HA expression in co-cultures

We next examined the effect of capsaicin on expression of Ca_V_2.2_HA at DRG presynaptic terminals, using RIM 1/2 expression as a presynaptic marker. In DIV 21 co-cultures, Ca_V_2.2_HA and RIM 1/2 labelling can be seen along processes on control and capsaicin-treated neurons (Fig. 5A). Enlargements from the Airyscan images (Fig. 5A) of individual rosette clusters of Ca_V_2.2_HA puncta that co-localized with RIM 1/2 show clear differences between the capsaicin-treated and the control conditions, after 60 min of rest (Fig. 5B). Analysis revealed that there was an increase in the relative distance between the centres of Ca_V_2.2_HA and RIM 1/2 in capsaicin-treated (0.49 ± 0.05 μm) compared to control (0.25 ± 0.03 μm) cocultures (Fig. 5Ci). Furthermore, at the same time point (60 min), there was a decrease in the overall percentage co-localization between Ca_V_2.2_HA and RIM 1/2 to 23.5 ± 2.9 % in capsaicin-treated cocultures, from 35.3 ± 1.2 % in controls (Fig. 5Cii). Although there was no significant increase in the relative distance between the centres of Ca_V_2.2_HA and RIM 1/2 at any other time point Supplementary Figs. 6A,B), a significant decrease in their percentage co-localization was also observed in capsaicin-treated neurons after 40 min (Supplementary Figs. 6A, C).

**Figure 5:**
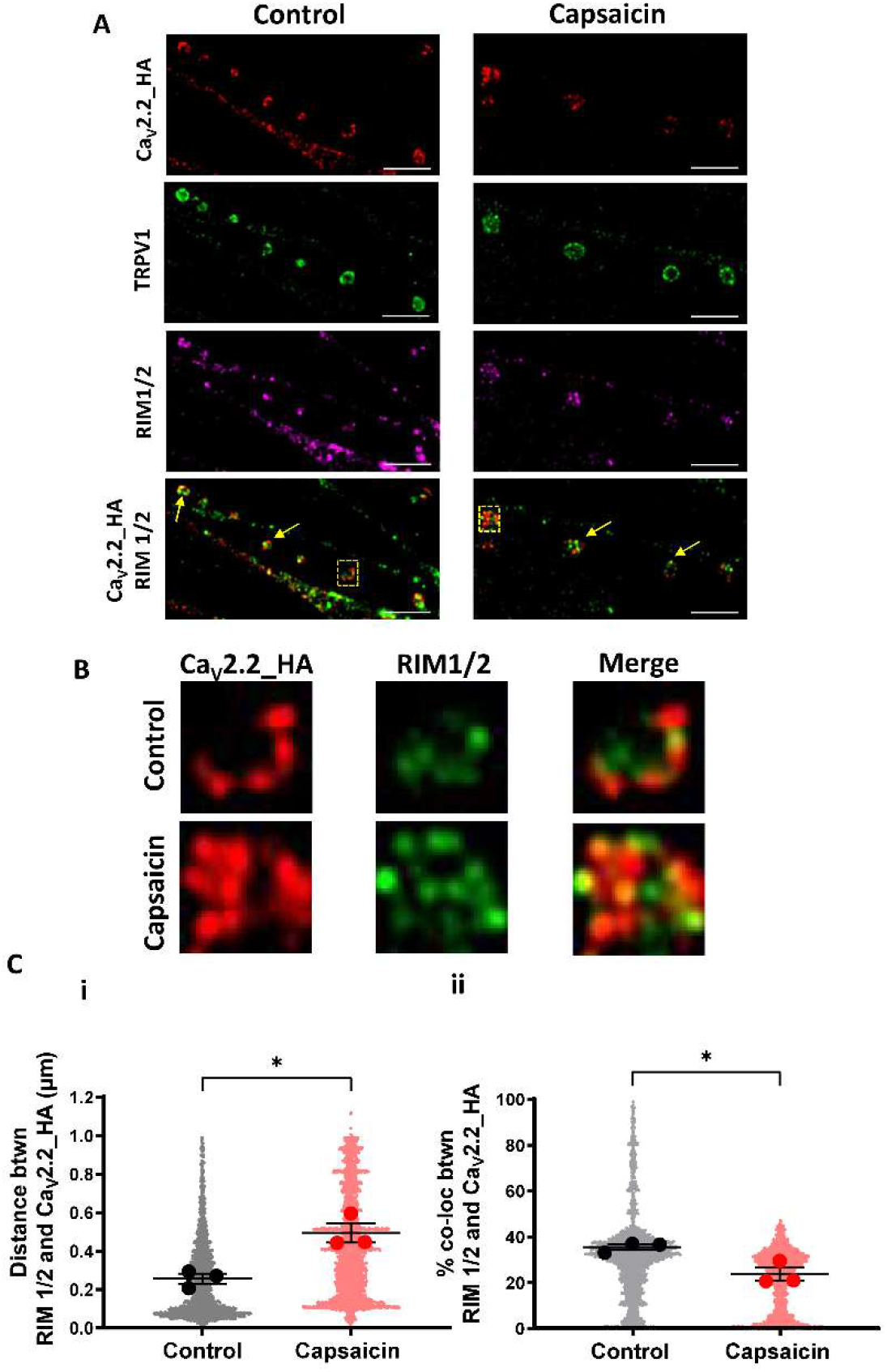
Decrease in Ca_V_2.2_HA immunolabelling in presynaptic terminals of DRG neurons following 2 min treatment with capsaicin. Capsaicin or control medium applied to DIV 21 co-cultures for 2 min, followed by 60 min rest at 37°C prior to fixation. (**A**) Images of presynaptic terminals from DRG neurons control (left) and capsaicin (right) conditions. Ca_V_2.2_HA (top row; red), TRPV1 (second row; green), RIM1/2 (third row; magenta), merged image of Ca_V_2.2_HA (red) and RIM 1/2 (re-coloured green for clarity; bottom row). Co-localized Ca_V_2.2_HA and RIM 1/2 marked by yellow arrows. Scale bars: 5 μm. (**B**) Images are enlargements of the ROI (2 x 2 μm dashed boxes) in (**A**), showing representative presynaptic terminals in control (top) and capsaicin (bottom) conditions. From left to right, Ca_V_2.2_HA (left; red), RIM 1/2 (middle; green) and merged panel (right). (**Ci**) Distance measurements between centres of co-localized objects Ca_V_2.2_HA and RIM 1/2. (**Cii**) Measurements of the percentage co-localizing volume for each object’s pair for Ca_V_2.2_HA and RIM 1/2. Individual data points represent distance and percentage measurements between colocalizing objects, 2300 pairs for Ca_V_2.2_HA and RIM 1/2, for both control and capsaicin conditions. Mean ± SEM of the three experiments are superimposed, with individual means as larger symbols. Statistical analysis: Student’s t-test * p<0.05.

### Capsaicin disturbs the contribution of the N-type calcium channel to Ca^2+^ transients

To determine the effect of capsaicin on the function of presynaptic N-type calcium channels, AP-mediated presynaptic Ca^2+^ elevation was examined using sy-GCaMP6f. Co-cultures were first stimulated with a 1 AP train followed by 200 APs at 10 Hz to determine the baseline responses of the synaptic boutons. Following a 5 min rest period, co-cultures were perfused with 1 μM capsaicin or control medium for 2 min and then perfused for 20, 40 or 60 min with control medium. To determine the contribution of N-type calcium channels, co-cultures were then re-stimulated with 1 AP in the presence or absence of CTX (Fig. 6A).

**Figure 6:**
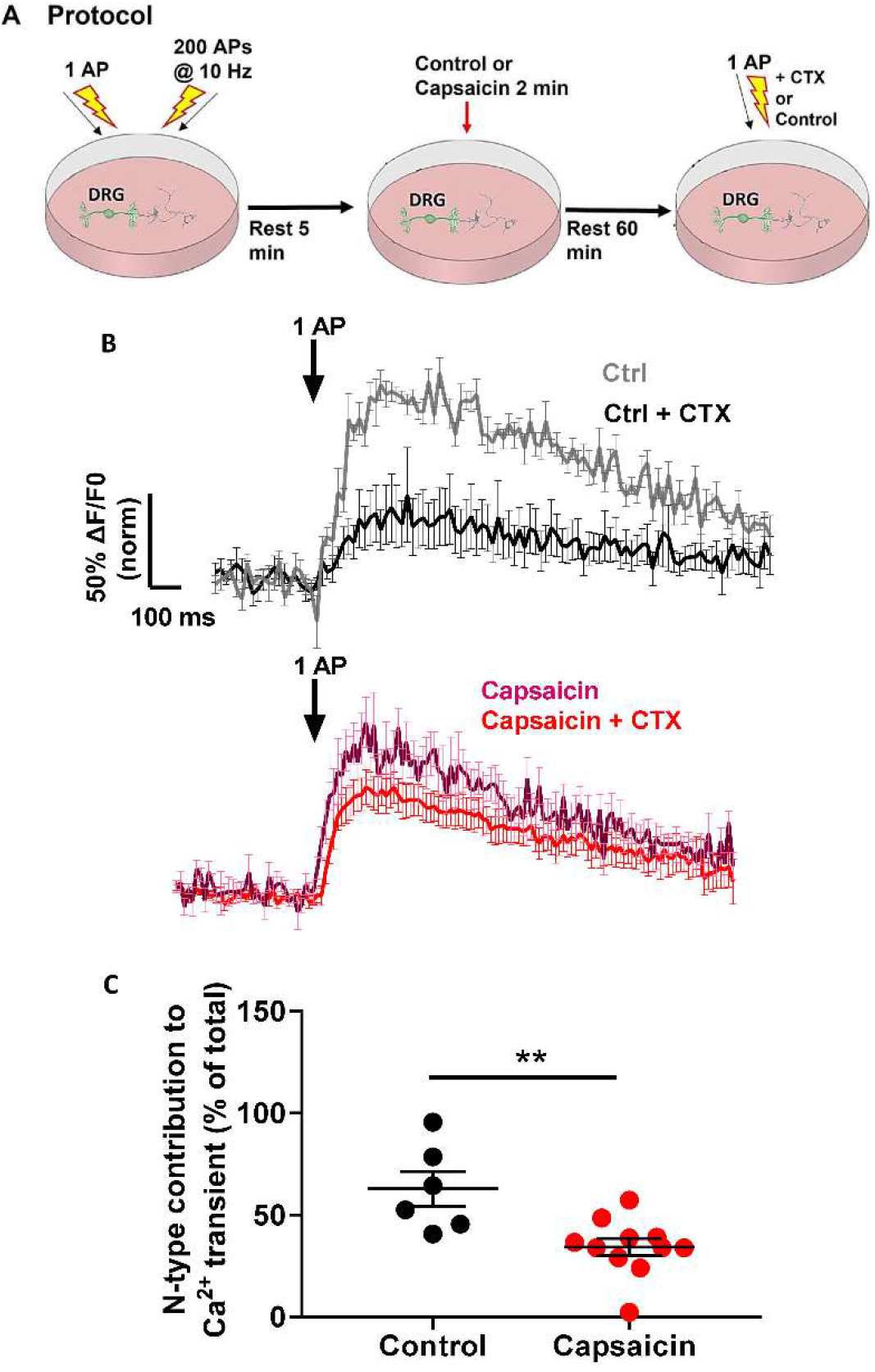
Reduction Ca_V_2.2 contribution to Ca^2+^ transients following 1 AP stimulation in capsaicin-treated, compared to control co-cultures following 60 min rest. (**A**) Schematic diagram of Ca^2+^ imaging protocol to test the effects of capsaicin in 21 DIV co-cultures. An initial protocol consisting of 1 AP and 200 APs (at 10 Hz) stimulations was applied to assess the baseline response of the neuron. Following a 5 min rest period, either control or capsaicin medium was applied for 2 min. Control medium was then applied for a 60 min rest period. CTX was then applied for 2 min. A final 1 AP stimulation was applied and the initial 1 AP Ca^2+^ transients were used to normalise the final 1 AP Ca^2+^ transients to determine the contribution of N-type calcium channels to the response. (**B**) Average normalised Sy-GCaMP6f fluorescence change in response to 1 AP stimulation, from control (top traces, pre-CTX grey (n = 8), post-CTX black (n = 8)) or capsaicin-treated (bottom traces, pre-CTX purple (n = 11), post-CTX red (n = 11)) neurons. The Ca^2+^ transients are expressed as ΔF/F0 and normalised to the averaged peak recorded from synaptic boutons before control medium/capsaicin and CTX was applied. Scale bars apply to both sets of traces. (**C**) Contribution of N-type voltage-gated calcium channels to the Ca^2+^ transient in response to 1AP. For Control + CTX (black circles) n = 6, and for capsaicin + CTX (red circles) n = 11. Statistical analysis: Mann-Whitney test; **p<0.01.

In control conditions, when neurons were perfused with control medium and allowed to rest for 60 min, we observed that CTX caused a large decrease of 63 % in the peak Ca^2+^ transient (control: 100 ± 6.0 %, n = 11 compared to control + CTX: 37.0 ± 3.0 %, n = 6) (Fig. 6B). However, when we compared the effects of CTX on capsaicin-treated co-cultures, it reduced the peak Ca^2+^ transient following a 1 AP stimulation by only 28 % (capsaicin: 80.0 ± 4.0 %, n = 8 compared to capsaicin + CTX: 57.0 ± 1.0 %, n = 11) (Figs. 6B). Thus there was a significant reduction in the effect of CTX following incubation with capsaicin on the 1 AP induced Ca^2+^ transient (Fig. 6C). A similar effect of capsaicin can be seen after 40 min but not after 20 min (Supplementary Fig. 7). Together, these data suggest that capsaicin reduces, with a slow timescale, the available N-type calcium channels that functionally contribute to Ca^2+^ transients following 1 AP stimulation.

### Reduced temperature inhibits capsaicin-induced loss of cell surface and active zone Ca_V_2.2_HA

We then examined whether endocytosis of Ca_V_2.2_HA from the cell surface of DRG neurons was contributing to the effect of capsaicin, by employing a reduced temperature (17°C), which inhibits trafficking processes such as endocytosis. Capsaicin was applied to co-cultures for 2 min at 37°C, followed by rest for 60 min at either 37 °C or 17°C. In small DRG neurons incubated at 37°C, a 37 % decrease in cell surface Ca_V_2.2_HA immunolabelling was observed (control: 1.0 ± 0.03; capsaicin: 0.63 ± 0.03) (Fig. 7A), similar to the results shown in Fig. 4D. In contrast, when co-cultures were incubated at 17°C, a significantly smaller decrease of 16 % in cell surface Ca_V_2.2_HA expression was measured (control: 1.0 ± 0.01; capsaicin: 0.84 ± 0.04) (Figs. 7Aii and 7B). Similarly, in medium DRG neurons, capsaicin treatment resulted in a smaller decrease in cell surface Ca_V_2.2_HA expression at 17°C compared to 37 °C (Supplementary Fig. 8).

**Figure 7:**
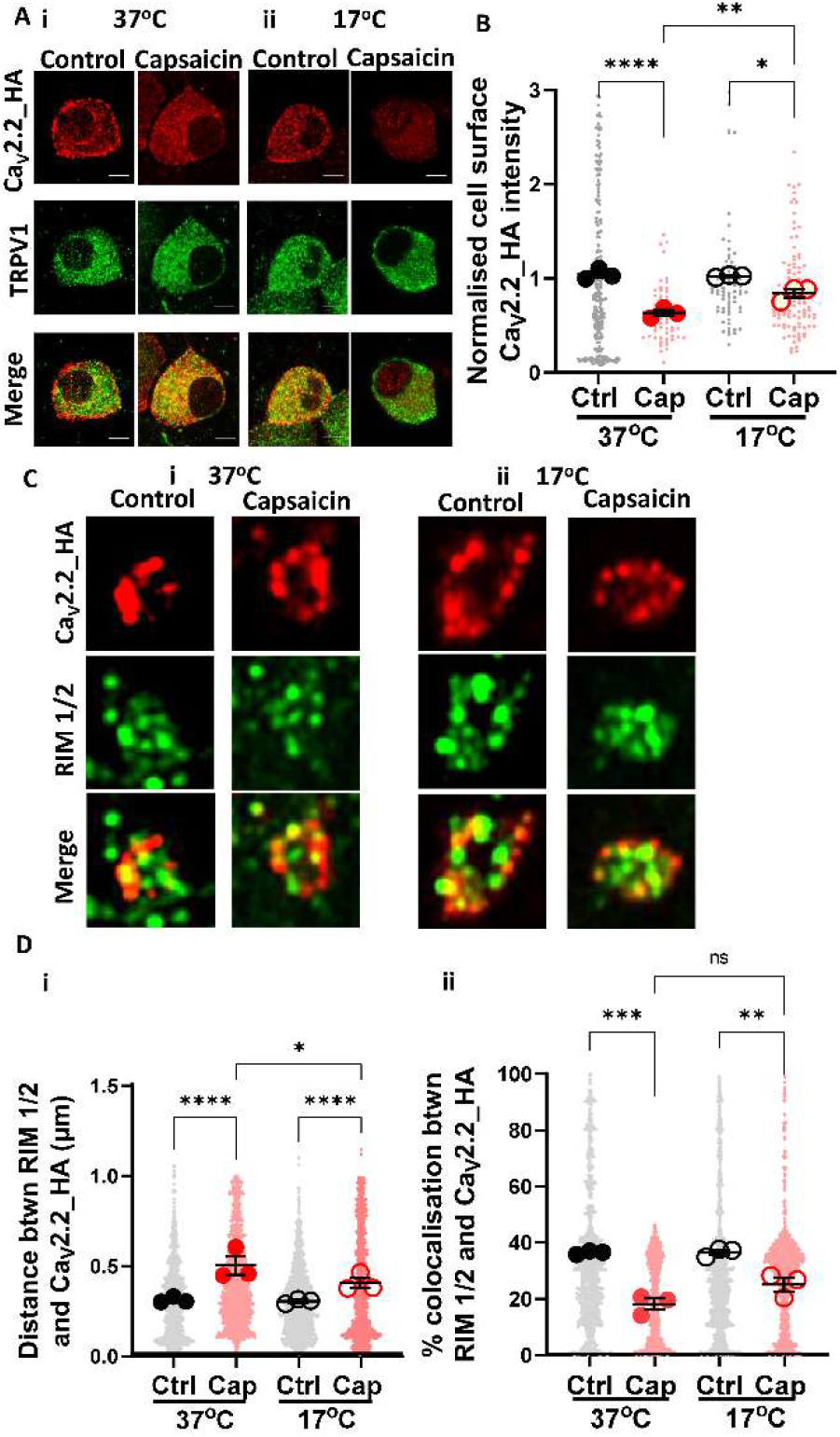
Decrease in cell surface Ca_V_2.2_HA in capsaicin-treated small DRG neurons and presynaptic terminals following incubation at 17°C. Capsaicin was applied to DIV 21 co-cultures for 2 min at 37°C, followed by 60 min rest at either 37°C or 17°C. (**A**) Images of small DRG neurons following (**i**) 37°C or (**ii**) 17°C rest period. Ca_V_2.2_HA (top; red), TRPV1 (middle; green) and merged panel (bottom). Scale bars: 5 μm. (**B**) Normalized cell surface Ca_V_2.2_HA intensity following 37°C (filled circles) or 17°C (open circles) rest. Individual data points represent Ca_V_2.2_HA intensity (normalized to mean in control conditions at 37°C), measured from three separate experiments and a total of 207 and 56 (control, grey) and 70, 106 (capsaicin, red) cells at 37°C and 17°C, respectively. Mean of each experiment shown by larger symbols. Mean ± SEM (n=3) is superimposed: (**C**) Images (2 x 2 μm ROIs) of control (left) and capsaicin-treated (right) terminals following incubation for 60 min at either (**i**) 37°C or (**ii**) 17°C. Ca_V_2.2_HA (top, red), RIM 1/2 (middle, green) and merged panel (bottom). (**Di**) Distance between centres of co-localized objects Ca_V_2.2_HA and RIM 1/2 in control and capsaicin conditions following rest at either 37°C or 17°C. (**Dii**) Percentage co-localizing volume for each object’s pair for Ca_V_2.2_HA and RIM 1/2 in control and capsaicin conditions following rest at either 37°C or 17°C. Data points represent distance and percentage measurements between individual co-localizing objects, 1500 pairs for Ca_V_2.2_HA and RIM 1/2, for control (black) and capsaicin (red) conditions at both temperatures. Mean of each experiment shown by larger symbols. Mean ± SEM (n = 3) superimposed. Statistical analysis: one-way ANOVA with Sidak’s multiple comparison post-hoc test; * p<0.05, **p<0.01, ***p<0.001, ****p<0.0001 and ns = not significant.

We next investigated the effect of capsaicin on expression of Ca_V_2.2_HA at presynaptic terminals following rest for 60 min at either 37°C or 17°C. At 37°C, there was a decrease in co-localization between Ca_V_2.2_HA and RIM 1/2 (Figs. 7C-D), as previously observed. The distance between Ca_V_2.2_HA and RIM 1/2 was significantly increased by 63 % from 0.30 ± 0.01 μm to 0.49 ± 0.05 μm (Fig. 7Di). In contrast, at 17°C the distance between the two centres increased to a significantly smaller extent, by 34 % (0.29 ± 0.01 μm to 0.39 ± 0.03 μm) (Figs. 7C, Di). However, there was a non-significant reduction in percentage colocalization of Ca_V_2.2_HA and RIM 1/2 due to capsaicin at 37°C (a 50.8 % decrease) and at 17°C (a 31.7 % decrease) (Fig. 7Dii). These data suggests that incubation at 17°C may impact Ca_V_2.2_HA distribution at both the cell body and presynaptic terminals by affecting a temperature-dependent process, such as endocytosis, induced by capsaicin.

### Dominant-negative Rab11a inhibits capsaicin-induced loss of functional Ca_V_2.2_HA in DRG terminals

We have previously shown that α2δ-1 and α2δ-2 are recycled to the plasma membrane via a Rab11a-dependent recycling endosome pathway ^11,12^. Furthermore, α2δ-1 increases Ca_V_2.2 at the plasma membrane by increasing the rate of the net forward trafficking of Ca_V_2.2 in a Rab11a-dependent manner ^12^. To test the involvement of Rab11a on the distribution of Ca_V_2.2_HA and the effect of capsaicin in cocultures, we transfected DRG neurons with dominant-negative Rab11a(S25N), as previously described for hippocampal neurons ^11,12^. We compared the effects of capsaicin on presynaptic Ca_V_2.2_HA expression in the absence and presence of Rab11a(S25N), using mCherry as a transfection marker (Fig. 8A). Similar Ca_V_2.2_HA immunolabelling was observed, apposed to RIM 1/2, at the presynaptic active zone when capsaicin was applied to co-cultures expressing either Rab11a(S25N) or empty vector (Fig. 8A). In control-transfected neurons expressing mCherry, we observed a 34 % decrease in Ca_V_2.2_HA immunostaining puncta associated with RIM 1/2 as a result of capsaicin treatment in the DRG terminals (Fig. 8B). In contrast, in DRG terminals expressing Rab11a(S25N), there was no significant difference in Ca_V_2.2_HA expression in the presence or absence of capsaicin (Fig. 8B). These data, supported by those in Fig. 7, provide evidence that presynaptic Ca_V_2.2_HA distribution is regulated by a Rab11a-dependent process, potentially involving capsaicin-induced endocytosis.

**Figure 8:**
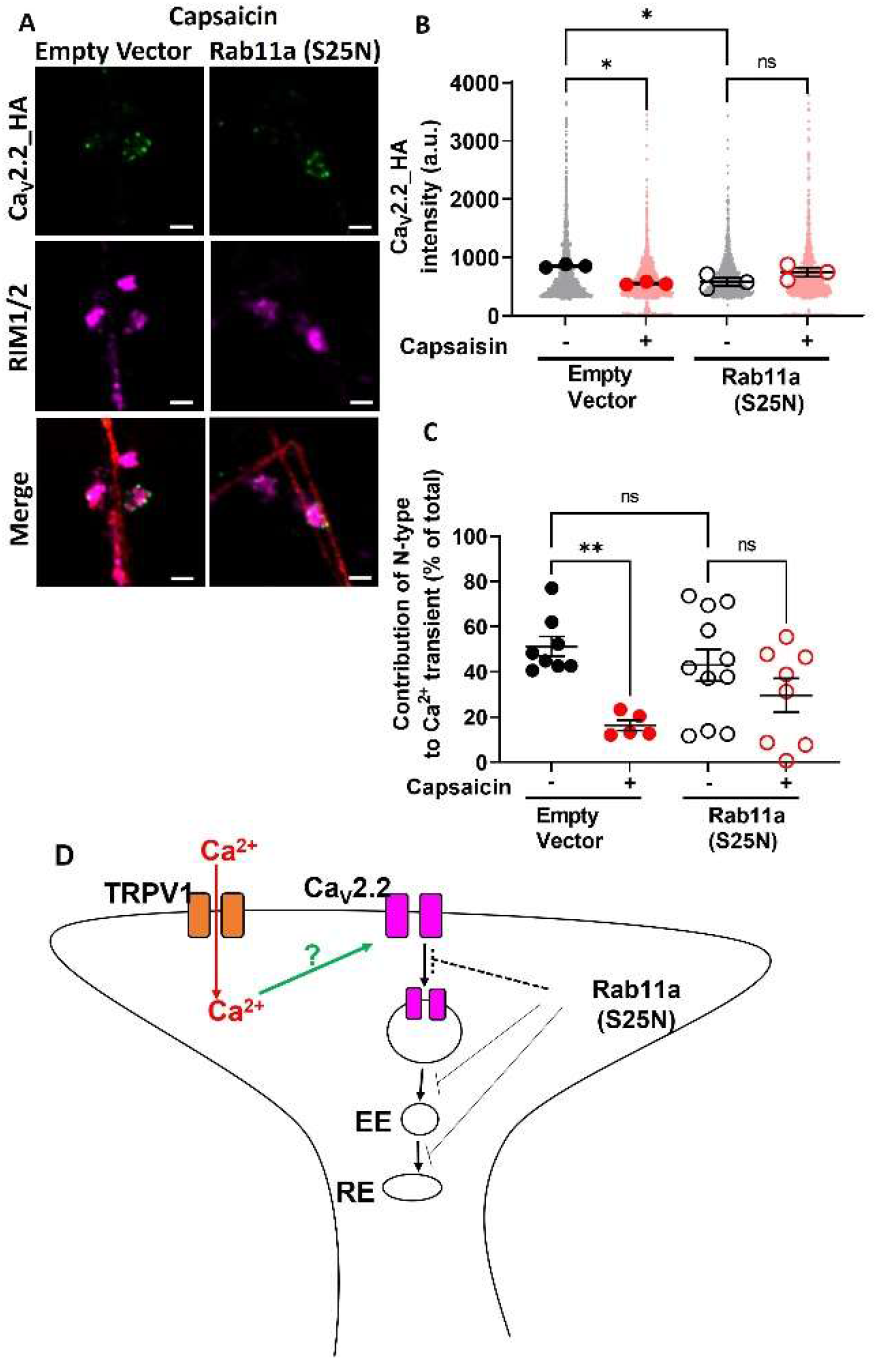
Effects of capsaicin on Ca_V_2.2_HA expression and presynaptic function are blocked by expression of dominant-negative Rab11a in co-cultured DRG neuron terminals. (**A**) Representative images showing capsaicin-treated co-cultures transfected with either empty vector (left) or Rab11a(S25N) (right). Ca_V_2.2_HA (green; top), RIM 1/2 (magenta; middle) and merged panel (bottom, including mCherry transfection marker). Scale bars 2 μm. (**B**) Mean Ca_V_2.2_HA intensity measured in presynaptic terminals (identified by presence of RIM 1/2) of DRG neurons transfected with either empty vector (left, solid symbols) or Rab11a(S25N) (right, open symbols) and treated with either control medium (black circles) or capsaicin (red circles). Individual data points represent 1400 puncta from all conditions, with means of each experiment in larger symbols. Mean ± SEM (n = 3) is superimposed. Statistical analysis: one-way ANOVA with Sidak’s multiple comparisons post-hoc test; * p<0.05 and ns = not significant. (**C**) Average normalised Sy-GCaMP6f fluorescence change in response to 1 AP stimulation recorded from synaptic boutons (DIV 21). Co-cultures were treated with either control medium (black circles) or capsaicin (red circles) followed by 60 min rest in control medium. Subsequently, CTX was applied, as in Fig. 6A. The Ca^2+^ transient is expressed as ΔF/F0 and normalised to the averaged initial peak in response prior to capsaicin. For Empty Vector transfected cells, control + CTX (n = 8; filled black circles) and capsaicin + CTX (n = 5; filled red circles), and for Rab11a(S25N) transfected cells, control + CTX (n = 11; open black circles) and Capsaicin + CTX (n = 8; open red circles). Statistical analysis: one-way ANOVA with Sidak’s multiple comparison post-hoc test; * p<0.05 and ns = not significant (**D**) Schematic diagram for Ca_V_2.2 membrane expression and recycling via Rab11a-positive endosomes. EE = early endosome, RE = recycling endosome. Dotted pathways represent hypothetical routes.

This result was supported by the measurement of AP-mediated presynaptic Ca^2+^ transients. The protocol outlined in Fig. 6A was applied to co-cultures of DRG neurons transfected with Rab11a(S25N) or control empty vector. Similar to the results seen in Figs. 6B, C, following the application of capsaicin, the N-type channel contribution to the 1 AP Ca^2+^ transient was markedly reduced from 51.3 ± 4.4 % (control) to 16.4 ± 2.3 % (capsaicin) for control-transfected DRGs (Fig. 8C). In contrast, in co-cultures expressing Rab11a(S25N), capsaicin produced no significant difference in the contribution of the N-type calcium channels to 1 AP-induced Ca^2+^ transients (control: 40.9 ± 8.2 %, compared to capsaicin: 29.4 ± 7.6 %). Together, these results suggest that interference with Rab11a-dependent function at presynaptic terminals may reduce endocytosis of Ca_V_2.2_HA in response to capsaicin.

## Discussion

### Ca_V_2.2_HA expression at the presynaptic boutons of DRG - spinal cord neuron co-cultures

Co-cultures of dissociated DRG and spinal cord neurons have been used as a model system to characterize the properties of primary afferent synapses ^21,22^. Importantly, DRG neurons do not form synapses between each other *in vivo* or in culture. In the present study, co-cultures were established by using DRG neurons from P0/P1 Ca_V_2.2_HA^KI/KI^ mice, cultured with spinal cord neurons from Ca_V_2.2^WT/WT^ mice. Thus, DRG neurons and their terminals could be distinguished based on the presence of Ca_V_2.2_HA and cell body size.

Our work first details the development of endogenous Ca_V_2.2_HA distribution during synapse formation between DIV 7 and 28, in in these co-cultures. DRG neuron terminals were detected using Ca_V_2.2_HA, together with presynaptic markers vGluT2 and RIM 1/2, together with the postsynaptic marker, Homer (Figs. 1 and 2). The predominant presynaptic boutons on central processes of primary afferents are of the *en passant* type, and have a diameter of 1 - 2 μm and length of 1 - 4 μm ^42,43^. In the co-culture system described, axonal boutons of similar sizes were observed.

We find a significant decrease in cell surface expression of Ca_V_2.2_HA at the cell body of small and medium DRG neurons in co-cultures over time (Supplementary Fig. 3). In parallel, there was a significant increase of Ca_V_2.2_HA expression was observed at DRG presynaptic terminals within the co-cultures (Fig. 1). After protein synthesis in neuronal somata, several mechanisms have been proposed to mediate the delivery of membrane proteins to axons. These include axonal transport within trafficking endosomes, and non-polarised delivery to the somatic membrane, followed by transcytosis and endosomal transport in axons ^44,45^. For example, it has been shown that trkA receptors are transported to axons from the soma by transcytosis during development of sympathetic neurons ^46^. The decrease in Ca_V_2.2_HA at the somatic plasma membrane (Supplementary Fig. 3) may be due to redirection of channels into axons of the developing neurons. Another method of presynaptic protein delivery is through local synthesis of proteins at axonal sites ^47^. However, further studies are required to determine whether any Ca_V_2.2 is synthesized locally in DRG terminals.

### Synaptic boutons of DRG - spinal cord neuron co-cultures form functional synapses

Native VGCCs have been classified in DRG neurons using specific pharmacological agents to elucidate their physiological contribution ^48–50^. DRG neurons have been shown to express N-type channels, as well as other calcium channels in differing proportions ^2,51^. In the present study, Ca^2+^ transients were recorded from co-cultures following a train of 1 AP stimuli (Fig. 3). In parallel with the changes in synaptic morphology, there was a corresponding increase in the number of functional synaptic boutons between DIV 7 and 28 (Fig. 3). To further dissect the contribution of Ca_V_2.2 channels to the Ca^2+^ transients, CTX was found to significantly reduce the amplitude of the 1 AP-induced Ca^2+^ transient, which is consistent with the previously described important role of N-type VGCCs in triggering glutamate release at primary afferent synapses ^52,53^. However, further studies are required to determine the contribution of other VGCCs to these Ca^2+^ transients.

### Capsaicin modulates presynaptic Ca_V_2.2_HA expression in co-cultures

The effects of capsaicin on VGCC expression in different cell types, and the mechanisms involved, remain unclear. Capsaicin has been found to indirectly reduce Ca^2+^ entry through VGCCs in rat trigeminal and hippocampal neurons ^54^, and gastric smooth muscle ^55^. Similarly, in rat sensory neurons, TRPV1 activation, through capsaicin, mediates an overall reduction in N-type calcium current ^20,56^. In contrast, in guinea pig DRG neurons, capsaicin shifts the VGCC current-voltage relationship to more hyperpolarized potentials, which would result in facilitated activation of VGCCs in response to smaller depolarization ^57^.

To understand the effect of capsaicin on Ca_V_2.2 channels, and its consequences for primary afferent transmission, we examined the impact of capsaicin on cell surface Ca_V_2.2_HA expression. We found firstly that cell surface Ca_V_2.2_HA was predominantly present on small and medium TRPV1-positive DRG neurons (Figs 4). Secondly, we found a significant reduction in cell surface Ca_V_2.2_HA in these TRPV1-positive neurons in response to brief incubation with capsaicin, which is detected 40 - 60 min later (Fig. 4 and Supplementary Figs. 4, 5). Previous studies have used a polyclonal antibody raised against an intracellular epitope in the II-III loop of Ca_V_2.2 to monitor potential internalization of the N-type calcium channel in rat DRG neurons in response to capsaicin ^20^. However, as this antibody cannot distinguish cell surface and intracellular Ca_V_2.2 expression, it is difficult to conclusively determine internalization of Ca_V_2.2 using such immunocytochemical methods.

Furthermore, in response to capsaicin we found a concomitant decrease in co-localization of Ca_V_2.2_HA with RIM 1/2 associated with presynaptic active zones (Fig. 5). In parallel, the N-type calcium channel contribution to 1 AP Ca^2+^ transients was decreased following capsaicin application (Fig. 6). The entry of Ca^2+^ through TRPV1 channels initiates a cascade of events within neurons. Wu et al. (2005) observed capsaicin-induced dephosphorylation of Ca_V_2.2 through Ca^2+^ -dependent activation of calcineurin in rat DRG neurons. The role of calcineurin in Ca^2+^-dependent regulation of Ca^2+^ influx has previously been examined in NG108-15 cells ^58^. These authors demonstrated a decrease in high-voltage activated (HVA) current when calcineurin was over-expressed, which was reversed by intracellular FK506 (calcineurin inhibitor), attesting to calcineurin-dependency. Furthermore, the effect was blocked by the Ca^2+^ chelator BAPTA, highlighting that it is a Ca^2+^-dependent mechanism ^58^.

### Dominant-negative Rab11a inhibits capsaicin-induced endocytosis of Ca_V_2.2_HA in DRG neurites

We next explored the involvement of endocytosis in the mechanism of action of capsaicin on presynaptic Ca_V_2.2_HA, by employing two strategies, reduced temperature (Fig. 7) and dominant-negative Rab11a (S25N) (Fig. 8). Rab11a belongs to the Rab family of small GTPases which are involved in many aspects of vesicular transport, via temporal and spatial interactions with multiple effectors ^59^, and Rab11 is required for the direct recycling of endosomes ^60^. It has been identified in many neuronal types as mainly residing in somatodendritic compartments ^61,62^, but it is also present in synaptic vesicles ^63^. The abrogation of Rab11a function by a dominant-negative form Rab11a (S25N); ^64^ has been shown to disrupt the ability of α2δ-1 to increase Ca_V_2.2 expression in hippocampal neurites ^12^. Furthermore, Rab11 facilitates activity-dependent bulk endocytosis ^65^, which suggests it has a vital role in neurotransmission during intense neuronal activity and Ca^2+^ influx. Rab11-dependent recycling of Ca_V_2.2 may therefore contribute to the dynamic control of expression of Ca_V_2.2-HA in presynaptic terminals of DRG neurons following TRPV1 activation.

Our data indicate that blockade of Rab11-dependent processes with Rab11a(S25N) leads to a reduction in cell surface Ca_V_2.2_HA levels that are associated with RIM 1/2 in presynaptic terminals, and capsaicin is no longer able to exhibit its down-regulatory effects on presynaptic Ca_V_2.2_HA (Figs. 8A, B). Furthermore, Rab11a(S25N) also prevented the capsaicin-induced reduction of the contribution of N-type calcium channel to 1 AP Ca^2+^ transients (Figs. 8C, D). Our results using a lowered incubation temperature of 17°C also point to the involvement of Ca_V_2.2_HA endocytosis in response to capsaicin (Fig. 7).

Previous work from our laboratory has shown that α2δ-1 and α2δ-2, but not α2δ-3 are recycled through Rab11a-dependent recycling endosomes and this process can be interrupted by gabapentin ^11,12^. Furthermore, in primary hippocampal neurites, Ca_V_2.2 membrane expression was found to be strongly dependent on the presence of an α2δ, and blockade of Rab11a-dependent recycling reduced cell surface Ca_V_2.2 levels in the presence of α2δ-1 ^12^. Our observations here extend these findings and show that expression of Rab11a(S25N) reduces levels of native Ca_V_2.2_HA in DRG neuronal presynaptic terminals Fig. 8B). It is also worth noting that Rab11-dependent recycling is an important mechanism by which cell surface expression of other ion channels is modulated, including K_V_1.5, KCNQ1 and epithelial TRPV5 channels ^66–68^.

The ability to examine plasma membrane expression and function of Ca_V_2.2 at presynaptic sites in the primary afferent pathway is critical for furthering our understanding of chronic pain and may suggest future routes for therapeutic targeting of this channel. Our results show that the use of DRGs from Ca_V_2.2_HA^KI/KI^ mice in co-culture with spinal cord neurons can be used to successfully examine the dynamic function and distribution of presynaptic Ca_V_2.2 channels and reveal the effect of TRPV1 activation on cell surface and presynaptic Ca_V_2.2_HA expression. Additionally, these data indicate that one of the main drivers of capsaicin-mediated decreases in Ca_V_2.2_HA from the plasma membrane involves a Rab11a-dependent process.

## Acknowledgements

This work was supported by a Wellcome Trust Investigator award to ACD (098360/Z/12/Z). We thank Dr. Manuela Nieto-Rostro and Wendy Pratt for advice during this project.

## Supplementary Figures

**Supplementary Figure 1:**
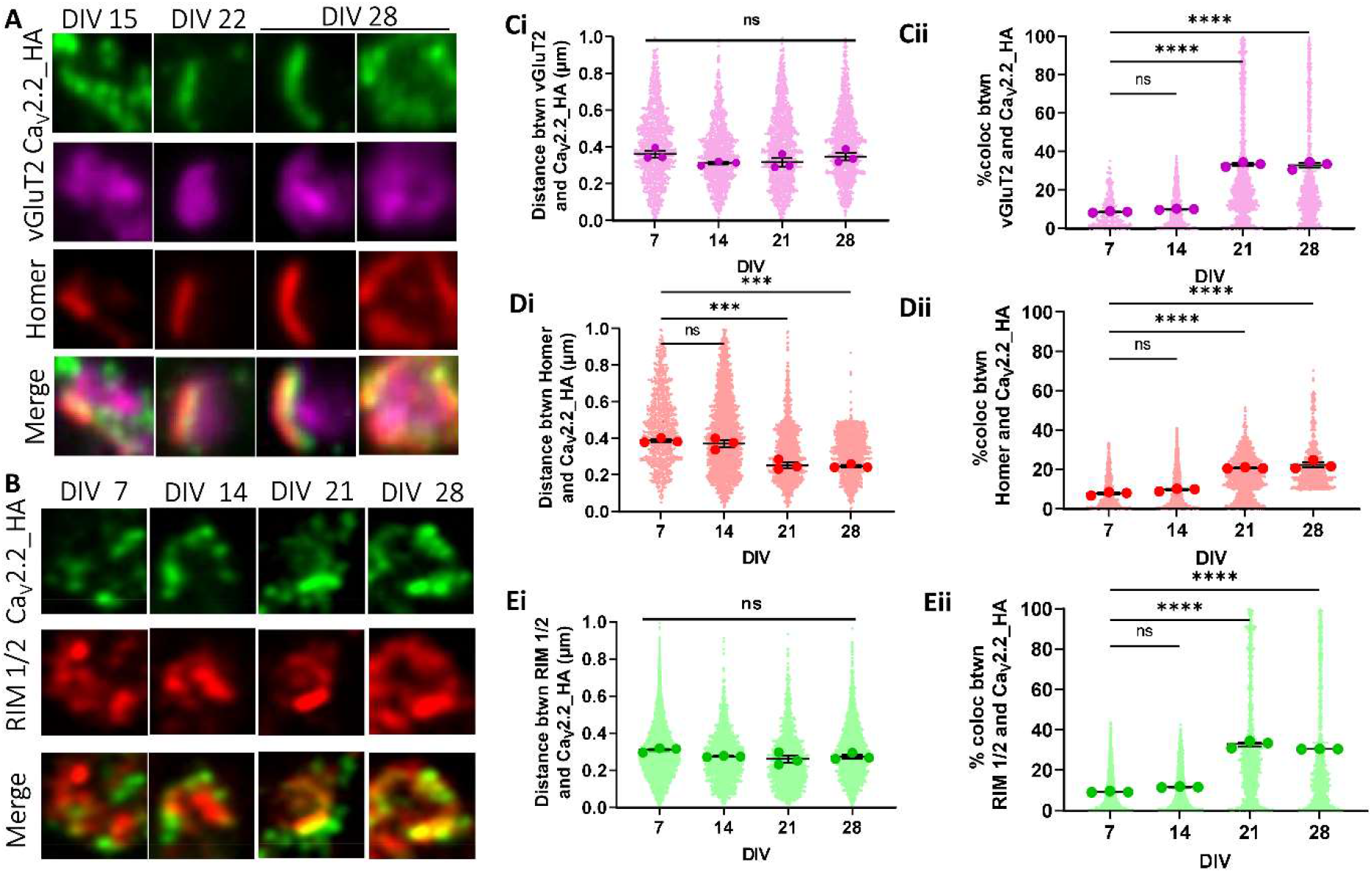
Increased co-localization of Ca_V_2.2_HA with vGluT2 and Homer at the presynaptic membrane in DRG - spinal cord co-cultures at DIV 21-28. (**A, B**) Representative images (2 x 2 μm) of synaptic puncta, (**A**) shows (top to bottom) Ca_V_2.2_HA (green), vGluT2 (magenta), Homer (red) and merged panels, fixed (left to right) at DIV 15, 22 and 28 (two images). (**B**) shows (top to bottom) Ca_V_2.2_HA (green), RIM 1/2 (red) and merged panels, fixed (left to right) at DIV 7, 14, 21 and 28. (**Ci-Ei**) Distance measurements between centres of co-localized objects for (**Ci**) Ca_V_2.2_HA and vGluT2, (**Di**) Ca_V_2.2_HA and Homer and (**Ei**) Ca_V_2.2_HA and RIM 1/2. (**Cii-Eii**) Measurements of the percentage co-localizing volume for each object’s pair for (**Cii**) Ca_V_2.2_HA and vGluT2, (**Dii**) Ca_V_2.2_HA and Home and (**Eii**) Ca_V_2.2_HA and RIM 1/2. Individual data points represent distance and percentage measurements between co-localizing objects, 1200, 2300 and 1400 for Ca_V_2.2_HA and vGluT2 and Ca_V_2.2_HA and Homer, respectively from DIV 7, 14-15, 21-22 and 28 for 3 separate experiments. Mean of each experiment is shown in larger symbols. Mean ± SEM (n = 3) is superimposed. Statistical analysis: one-way ANOVA with Tukey selected comparisons post hoc test; ****p<0.0001, ***p<0.001, **p<0.01, *p<0.5, ns = not significant.

**Supplementary Figure 2:**
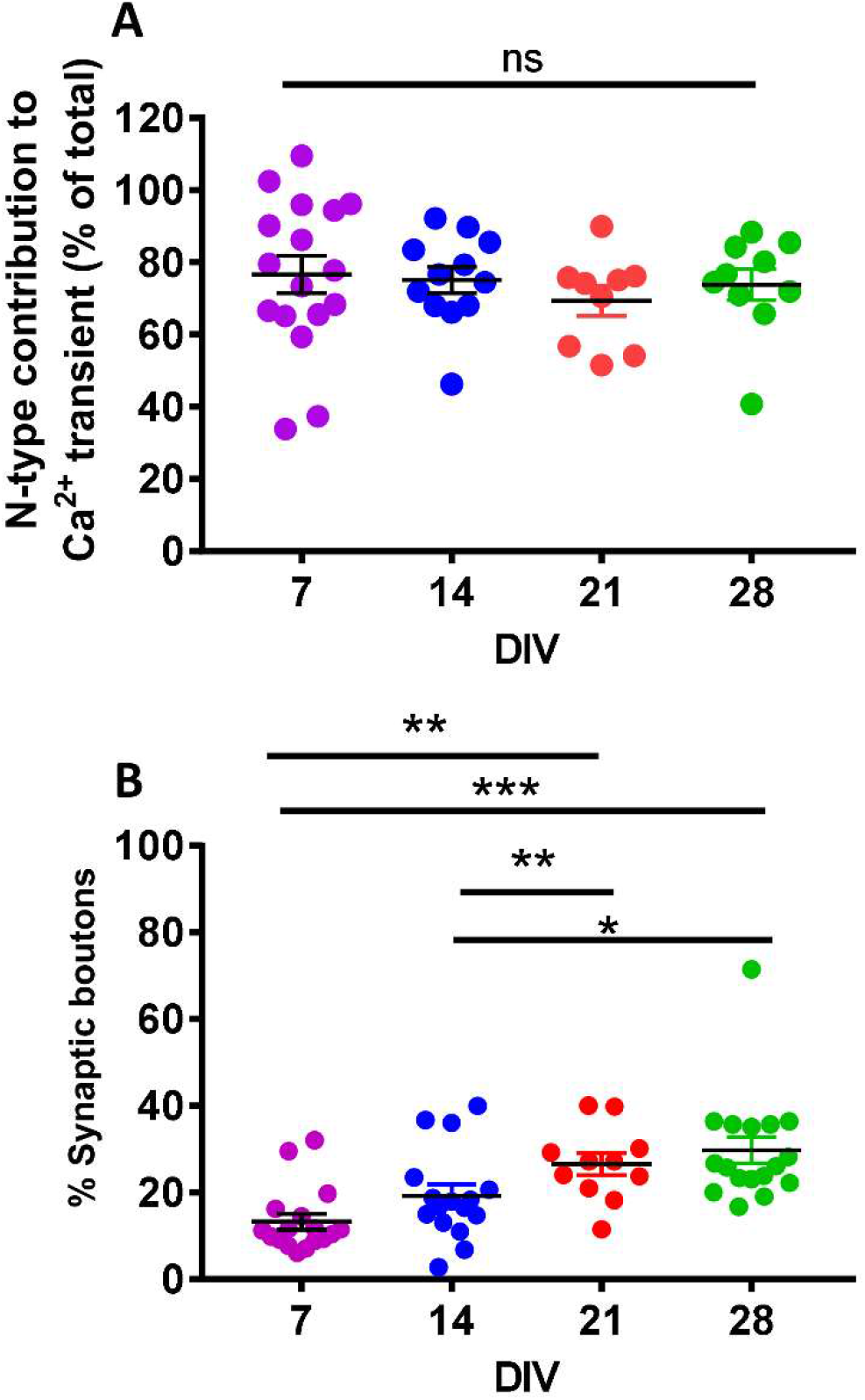
Properties of presynaptic 1 AP Ca^2+^ transients in DRG neuron terminals from co-cultures using Sy-GCaMP6f and VAMP-mOr2. (**A**) CTX was used to determine the N-type VGCC contribution to Ca^2+^ transients in response to 1 AP stimulation of co-cultures at increasing DIV. N-type VGCC contribution was 76.6 ± 5.1 % (n = 17), 75.1 ± 3.6 % (n = 12), 69.3 ± 4.2 % (n = 9) and 73.9 ± 4.3 % (n = 10), for DIV 7 (magenta), 14 (blue), 21 (red) and 28 (green), respectively. (**B**) % of VAMP-positive boutons at DIV 7 (n = 17; magenta), 14 (n = 13; blue), 21 (n = 11; red) and 28 (n = 17; green). Statistical analysis for (**A**) and (**B**): one-way ANOVA with Bonferroni selected comparison post hoc test; ***p<0.0001, ** p<0.001, *p<0.05; ns, not significant.

**Supplementary Figure 3:**
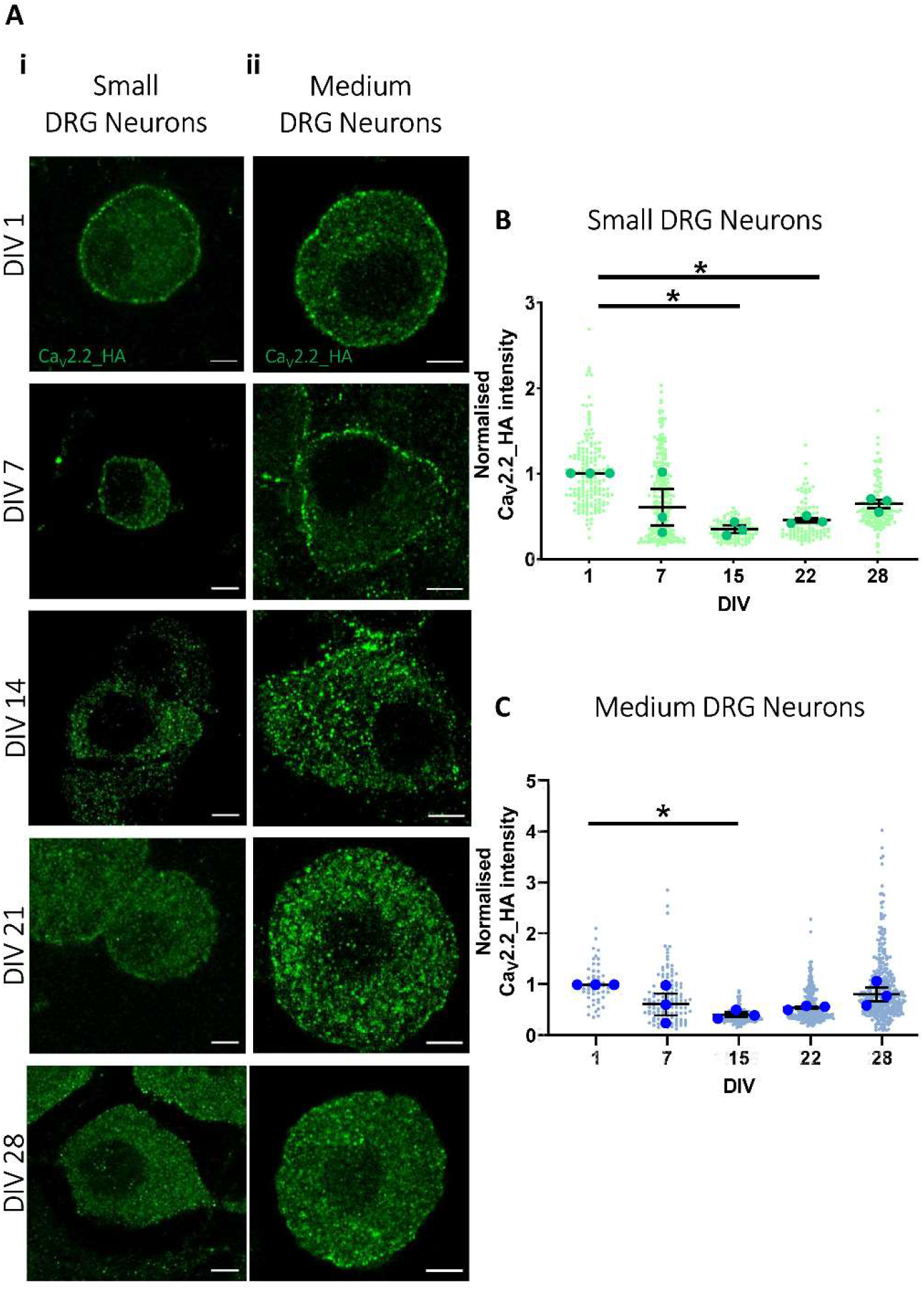
Cell surface Ca_V_2.2_HA expression on DRG cell bodies in co-cultures decreases over time *in vitro*. (**Ai**) Small and (**Aii**) medium DRG neurons from co-cultures composed of DRG neurons from Ca_V_2.2_HA^KI/KI^ and spinal cord neurons from Ca_V_2.2^WT/WT^ mice. Airyscan images of Ca_V_2.2_HA (green) at (top to bottom) DIV 1, 7, 14, 21 and 28. Scale bars for small and medium DRG neurons: 10 μm and 15 μm, respectively. (**B**) and (**C**) Cell surface Ca_V_2.2_HA intensity was measured from all Ca_V_2.2_HA-positive neurons at DIV 1. The mean fluorescence intensity values were then used to normalise cell surface Ca_V_2.2_HA of individual neurons depending on size, in 3 separate experiments, for (**B**) small and (**C**) medium DRG neurons at DIV 1, 7, 14, 21 and 28. Individual data points are shown for a total of 167, 203, 98, 103 and 122 small DRG neurons (**B,**green symbols) and 83, 119, 164, 262 and 359 medium DRG neurons (**C**, blue symbols), respectively. The mean for each experiment is shown in larger symbols and mean ± SEM (n=3) is superimposed. Statistical analysis: one-way ANOVA with Tukey selected comparison post-hoc test; * p<0.05.

**Supplementary Figure 4:**
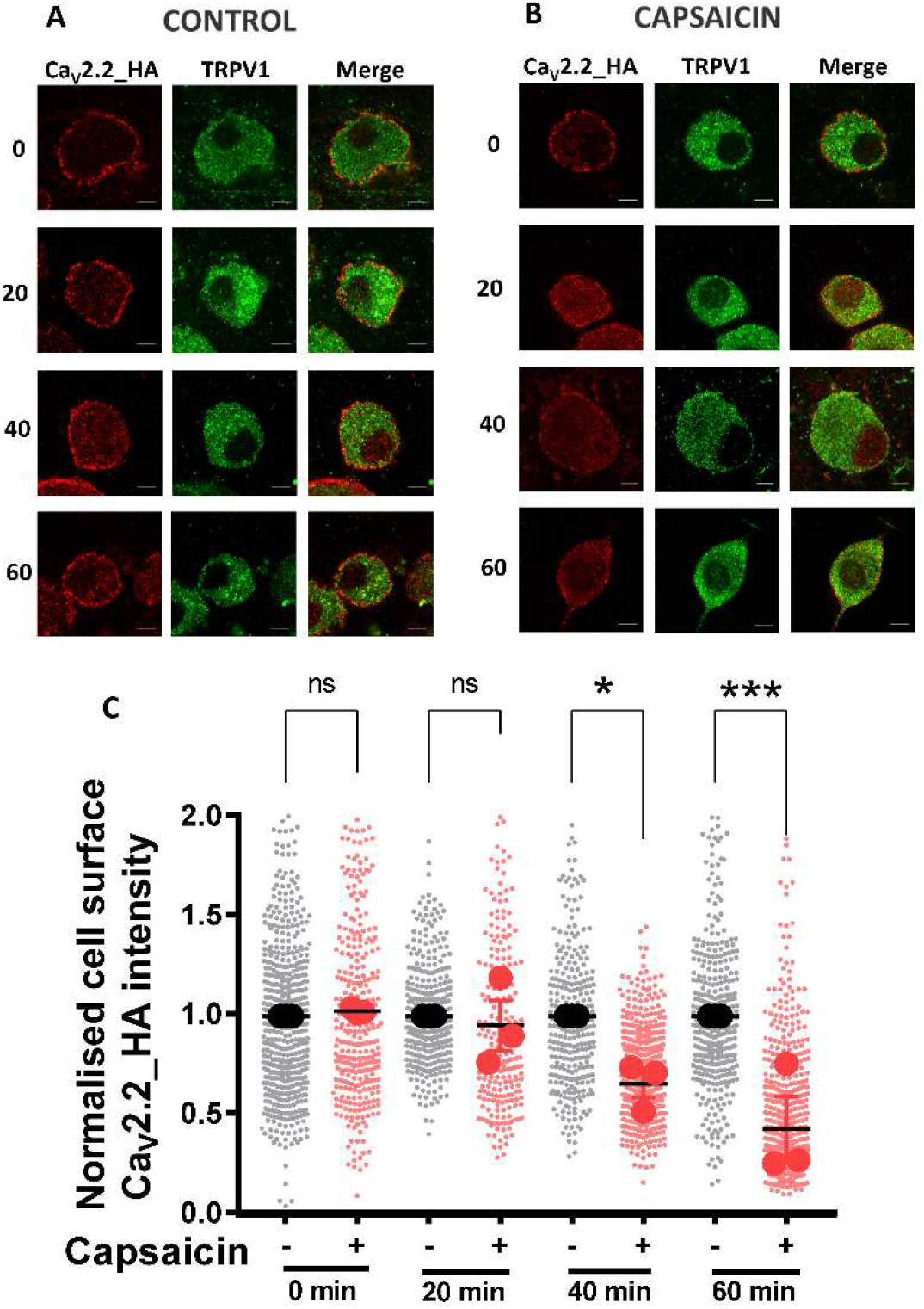
Decrease in cell surface Ca_V_2.2_HA in small DRG neurons at 40 and 60 min after application of capsaicin. Capsaicin or control medium (2 min) was applied to DIV 21 co-cultures, followed by rest for 0, 20, 40 and 60 min at 37°C prior to fixation. Representative images of small DRG neurons from control (**A**) and capsaicin (**B**) conditions. Ca_V_2.2_HA (left, red), TRPV1 (middle, green) and merged image (right), after rest for (top to bottom) and 0, 20, 40 and 60 min. Scale bar: 5 μm. (**C**) Normalized cell surface Ca_V_2.2_HA intensity following control or capsaicin application and rest for 0, 20, 40 and 60 min. For analysis, cell surface Ca_V_2.2_HA fluorescence intensity was measured from all TRPV1-positive neurons and the perimeter of these DRG neurons was used as an estimation of neuron size, the size-groups were analysed independently. In all experiments, the fluorescence intensity of each neuron was normalised against the mean fluorescence of control DRG neurons from their respective size-groups. Individual data points represent normalized Ca_V_2.2_HA intensity measured from three separate experiments and a total of 554, 354, 297 and 357 (control, grey circles) and 317, 249, 542 and 548 (capsaicin, red circles), respectively, normalised to control timepoint 0 min. Mean values for each experiment are shown by larger symbols, and mean ± SEM (n=3) is superimposed. Statistical analysis: one-way ANOVA with Sidak’s multiple comparisons post-hoc test; * p<0.05; ***p<0.001; ns, not significant.

**Supplementary Figure 5:**
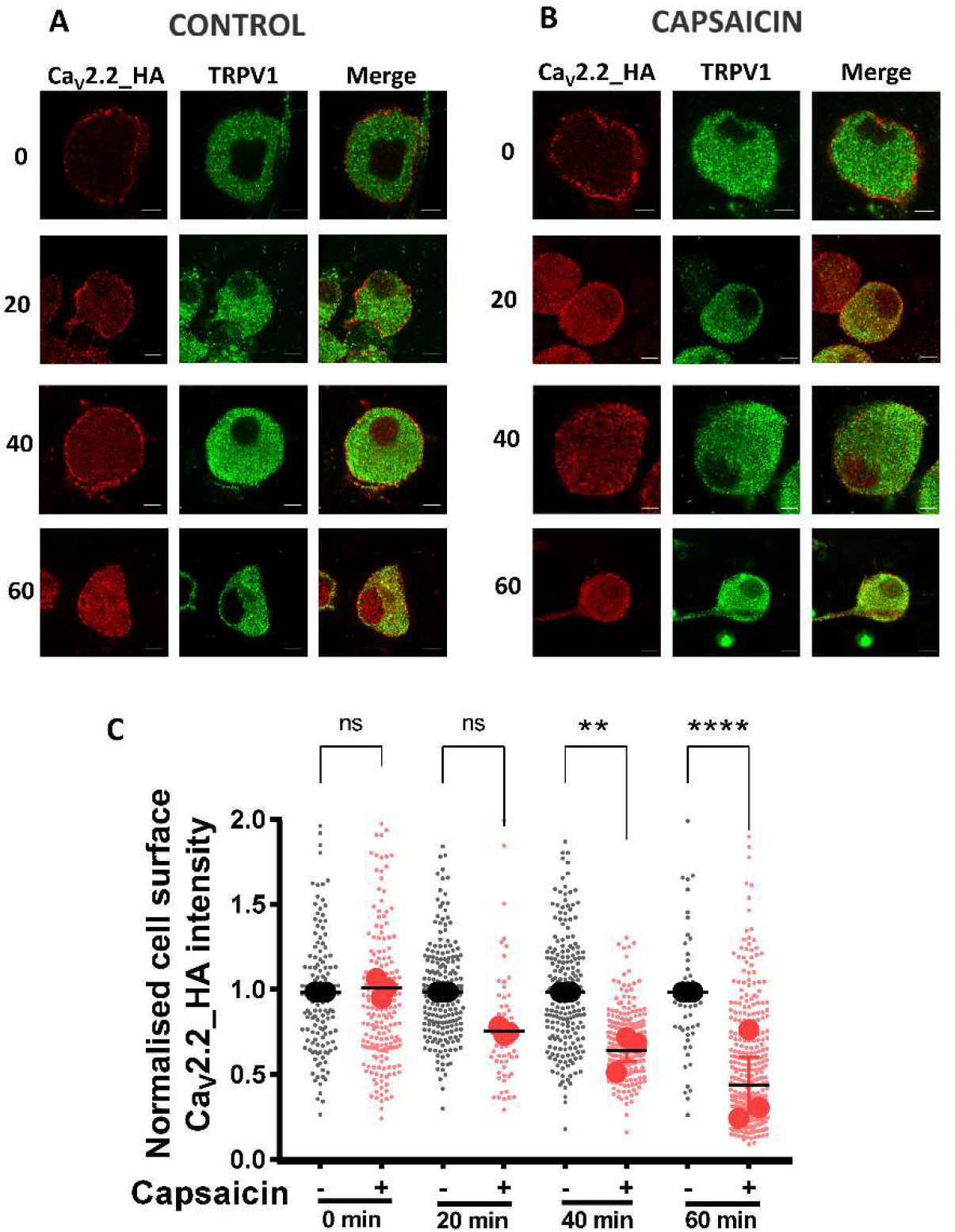
Decrease in cell surface Ca_V_2.2_HA in medium DRG neurons at 40 and 60 min after application of capsaicin. Capsaicin or control medium (2 min) was applied to DIV 21 co-cultures, followed by rest for 0, 20, 40 and 60 min at 37°C prior to fixation. Representative images of medium DRG neurons from control (**A**) and capsaicin (**B**) conditions. Ca_V_2.2_HA (left, red), TRPV1 (middle, green) and merged image (right), after rest for (top to bottom) and 0, 20, 40 and 60 min. Scale bar: 5 μm. (**C**) Analysis was performed as described in the legend to Supplementary Fig. 5. Normalized cell surface Ca_V_2.2_HA intensity following control or capsaicin application and rest for 0, 20, 40 and 60 min. Individual data points represent normalized Ca_V_2.2_HA intensity measured from three separate experiments, and a total of 140, 209, 210 and 50 (control, grey circles) and 199, 64, 229 and 343 (capsaicin, red circles), respectively, normalised to control timepoint 0 min. Mean values for each experiment are shown by larger symbols, and mean ± SEM (n=3) is superimposed. Statistical analysis: one-way ANOVA with Sidak’s multiple comparisons post-hoc test; ** p<0.01; **** p<0.0001; ns, not significant.

**Supplementary Figure 6:**
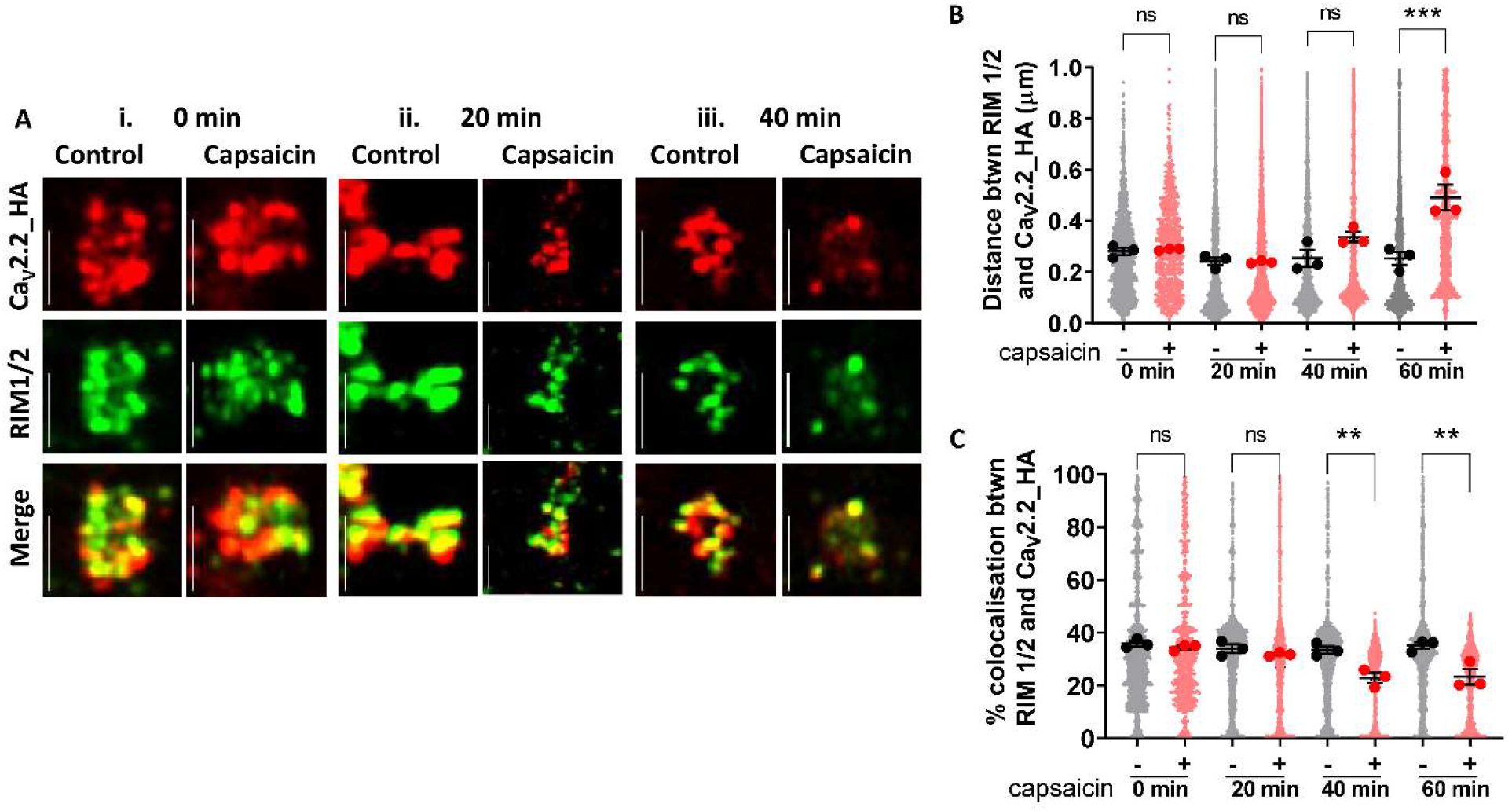
Ca_V_2.2_HA co-localization with RIM 1/2 is reduced with a slow time course in presynaptic terminals of DRG neurons following application of capsaicin. (**A**) Representative images of DRG presynaptic terminals following capsaicin (2 min application), then rest for (**i**) 0, (**ii**) 20, (**iii**) 40 min at 37°C prior to fixation. Ca_V_2.2_HA (red, left), RIM 1/2 (green, middle) and merged image (right) in control and capsaicin conditions. Scale bars: 2 μm. (**B**) Distance measurements between centres of co-localized objects Ca_V_2.2_HA and RIM 1/2. (**C**) Measurements of the percentage co-localizing volume for each object’s pair for Ca_V_2.2_HA and RIM 1/2. For (**B**) and (**C**), individual data points represent distance and percentage measurements between co-localizing objects for 2300 pairs, for both control (black) and capsaicin (red) conditions. Mean data for each of 3 experiments is shown in larger symbols and mean ± SEM (n = 3) is superimposed. Statistical analysis: oneway ANOVA with Sidak’s multiple comparisons test as post-hoc test; **p<0.01, ***p<0.001, ns = not significant.

**Supplementary Figure 7:**
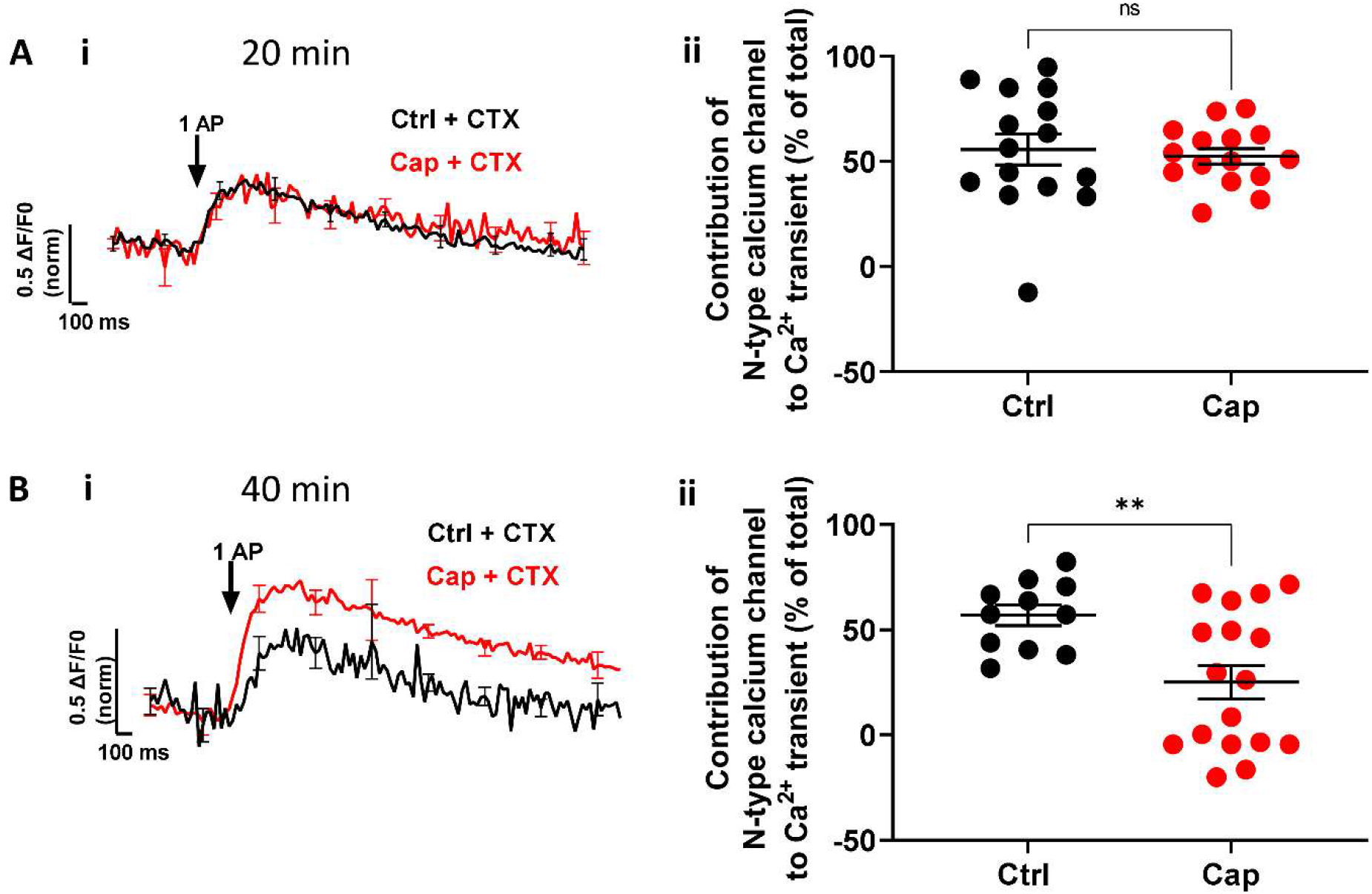
Capsaicin application reduces N-type VGCC contribution to 1 AP Ca^2+^ transients following 20 or 40 min rest. (**A i**) and (**B i**) Average normalised Sy-GCaMP6f fluorescence change in response to 1 AP stimulation recorded from VAMP positive DRG synaptic terminals of capsaicin (2 min)-treated or control DIV 21 co-cultures, followed by (**A**) 20 or (**B**) 40 min rest. Control medium (black trace) or capsaicin (Cap; red trace), followed by rest for time stated, then CTX application (according to protocol in Fig. 6A). (**A ii**) and (**B ii**) CTX was used to determine the N-type VGCC contribution to 1 AP stimulation. The 1 AP Ca^2+^ transient is expressed as ΔF/F0 and normalised to the averaged peak recorded from VAMP-positive boutons before control medium or capsaicin and CTX was applied. For (**A ii**) control (black circles) n = 15 and Capcaisin (red circles) n = 15. For (B ii) control (black circles) n = 11 and capsaicin (red circles) n = 17. Mean ± SEM are superimposed. Statistical analysis: Mann-Whitney test; *p<0.05, ns = not significant.

**Supplementary Figure 8:**
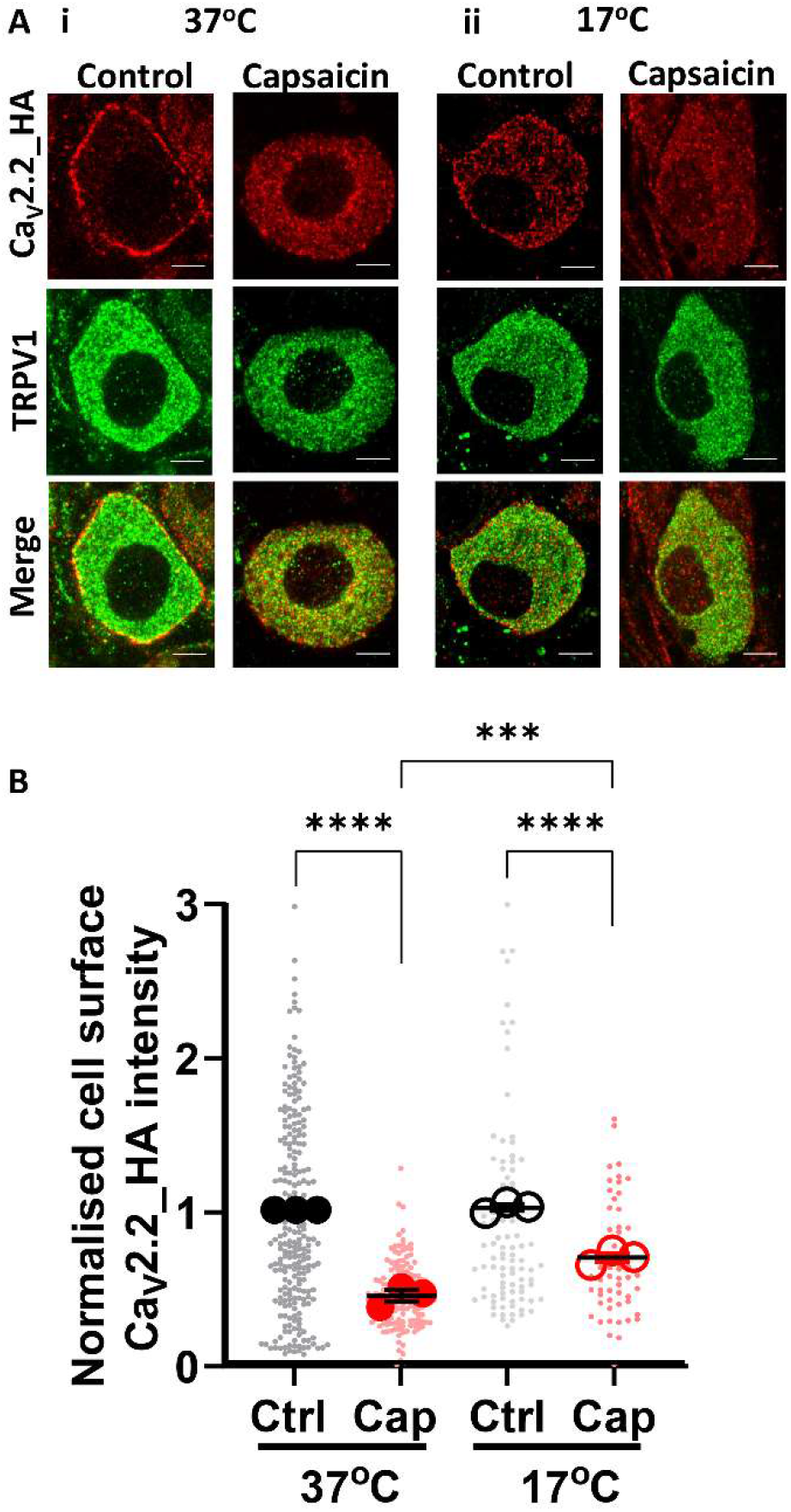
Decrease in cell surface Ca_V_2.2_HA in capsaicin-treated medium DRG neurons following rest at 17°C. (**A**) Capsaicin or control medium (2 min at 37°C) application to co-cultures (DIV 21), was followed by rest for 60 min at (**i**) 37°C or (**ii**) 17°C, prior to fixation. Representative images from medium DRG neurons showing Ca_V_2.2_HA (top, red), TRPV1 (middle, green) and merged image (bottom). Scale bars: 5 μm. (**B)**Ca_V_2.2_HA intensity (individual data points are normalized to mean value of medium DRG neurons in control conditions at 37°C). Individual data points measured from three separate experiments and a total of 235 and 154 cells for control and capsaicin conditions at 37°C (grey and red solid circles, respectively) and 101 and 54 from control and capsaicin conditions at 17°C (grey and red open circles, respectively). Mean values for each experiment are shown by larger back and red symbols. Mean ± SEM (n=3) is superimposed. Statistical analysis: one-way ANOVA with Sidak’s multiple comparisons post-hoc test; ***p<0.001 and ****p<0.001.

